# Genomic analysis reveals HDAC1 regulates clinically relevant transcriptional programs in pancreatic cancer

**DOI:** 10.1101/2022.09.06.506214

**Authors:** Carter A. Wright, Emily R. Gordon, Sara J. Cooper

**Affiliations:** The University of Alabama in Huntsville, Huntsville, AL, 35899, USA; HudsonAlpha Institute for Biotechnology, Huntsville, AL, 35806, USA

## Abstract

Novel strategies are needed to combat multidrug resistance in pancreatic ductal adenocarcinoma (PDAC). We applied genomic approaches to understand mechanisms of resistance in order to better inform treatment and precision medicine. Altered function of chromatin remodeling complexes contribute to chemoresistance. Our study generates and analyzes genomic and biochemical data from PDAC cells overexpressing *HDAC1*, a histone deacetylase involved in several chromatin remodeling complexes. We characterized the impact of overexpression on drug response, gene expression, HDAC1 binding, and chromatin structure using RNA-sequencing and ChIP-sequencing for HDAC1 and H3K27 acetylation. Integrative genomic analysis shows that *HDAC1* overexpression promotes activation of key resistance pathways including epithelial to mesenchymal transition, cell cycle, and apoptosis through global chromatin remodeling. Target genes are similarly altered in patient tissues and show correlation with patient survival. We also demonstrate that direct targets of HDAC1 that also show altered chromatin are enriched near genes associated with altered GTPase activity. HDAC1 target genes identified using *in vitro* methods and observed in patient tissues were used to develop a clinically relevant nine-transcript signature associated with patient prognosis. Integration of multiple genomic and biochemical data types enables understanding of multidrug resistance and tumorigenesis in PDAC, a disease in desperate need of novel treatment strategies.

## INTRODUCTION

Pancreatic ductal adenocarcinoma (PDAC) is the most common form of pancreatic cancer and one of the most lethal cancers with a five-year survival rate of 11.5%. Due to the lack of early-stage symptoms, 52% of patients are diagnosed with unresectable, locally advanced, metastatic cancer^1^. Chemotherapeutics such as gemcitabine, abraxane, and combination therapies like FOLFIRINOX are the standard of care, but chemoresistance develops in the majority of patients and contributes to the poor outcomes.

Several HDAC inhibitors (HDACi) have been tested (e.g. entinostat, romidepsin) for the treatment of solid tumors and hematological malignancies^2^. Four HDACi are FDA approved for treatment of hematological neoplasms (i.e., T-cell lymphomas and multiple myeloma)^3^’^4^. However, clinical trials for HDACis in PDAC have been largely unsuccessful^2^. HDAC inhibition by commercially available HDACis leads to negative side effects in patients^4^ since HDACs have a broad impact on expression of genes involved in cancer pathways and normal cellular functions including the function of non-histone proteins; thus, inhibition of multiple HDACs with molecules typically targeting an entire class of HDACs causes massive disruption to cellular function that has limited their clinical usage in treating solid tumors.

PDAC tumor cells achieve drug resistance through many cellular mechanisms: dysregulation of drug transporters, increased metabolism of drugs, upregulation of DNA repair, alterations in cell cycle and evasion of apoptosis Resistance is also associated with the epithelial to mesenchymal transition (EMT), the acquisition of a more mesenchymal cell state that is more invasive and has increased migratory potential^5,6^. While the exact mechanisms linking EMT to resistance are not well-described they are regulated by key transcription factors including SNAI1/2, ZEB1/2, and TWIST1/2. Additionally, the hypovascular nature of PDAC tumors prevents sufficient delivery of oxygen and drugs to the tumor cells, further exacerbating chemoresistance. This hypoxic environment promotes expression of anti-apoptotic genes (i.e. *STAT3, CDK6, CDK17, CDKN1A*), suppressing apoptosis and also contributing to drug resistance^5,7–9^. Drug resistance mechanisms often contribute to resistance to multiple drugs which makes the understanding of this complex problem challenging. Understanding mechanisms of drug resistance is necessary to facilitate development of therapeutic strategies to prevent or reverse resistance.

Previous work, including our own publications, links the expression of chromatin remodeling genes to chemoresistance and patient survival in PDAC^10–12^. Chromatin remodeling is a mechanism of gene regulation through rearrangement of chromatin structure to alter DNA accessibility and influence transcription factor binding. This process can alter gene expression patterns and lead to cellular reprogramming that contributes to chemoresistance. Dysregulation of chromatin remodeling genes leads to global changes in gene expression making it difficult to determine which subset of genes are most important for resistance, especially when many pathways are known to be involved^13^. Three key complexes involved in chromatin remodeling are the NuRD, Sin3A and CoREST complexes and they all include the key histone modifying protein, HDAC1^14^.

*We* previously demonstrated that overexpression of *HDAC1* contributes to multidrug resistance in pancreatic cancer cells^15^. While HDACs are canonically members of repressive complexes, binding of HDAC1 has also been associated with transcriptional activation. In some cases, this is explained by HDAC1’s ability to recruit RNA Pol II or regulate transcriptional elongation^16,17^. HDAC1 regulates the acetylation of histone and non-histone proteins to modulate gene expression and its overexpression has been associated with progression, metastasis, and patient prognosis in many cancer types including gastric, breast, colon, and prostate cancers^13^. In colorectal cancer HDAC1 promotes tumorigenesis by regulating the HIF1α/VEGFA signaling pathway via post-transcriptional modulation^18^. Just a few examples of the broad influences of HDAC1 include its recruitment to the promoter of *CDH1*, an epithelial cell marker, where it silences *CDH1* expression during metastasis^13^. HDAC1 also regulates expression of genes involved in resistance pathways in many cancers including apoptosis, DNA damage repair, metastasis, and EMT^13,19^. It was previously shown that HDAC1 is essential for cell proliferation and the transcription of core regulatory transcription factors (TFs), an essential factor in cancer growth. Due to the autoregulating nature of these TFs, HDAC inhibition led to their depletion through disruption of chromatin architecture and antiproliferative effects^20^. HDAC1’s regulation of genes involved in key cancer pathways including drug resistance, cancer progression, and tumor suppression make it a strong candidate as a drug target.

In this study, we characterized the impacts of *HDAC1* overexpression in a well described PDAC cell line, MIA PaCa-2, by measuring global effects on gene expression using RNA-sequencing, HDAC1 binding using ChIP-sequencing, and chromatin structure using H3K27ac histone profiling to better understand how *HDAC1* overexpression impacts key aspects of tumorigenesis and relevance for patient treatment. We found that *HDAC1* overexpression alters activity of several pathways (e.g. EMT, resistance to apoptosis, altered cell cycle checkpoint, and increased hypoxia) known to contribute to drug resistance. We showed that *HDAC1* overexpression leads to a more mesenchymal phenotype *in vitro* and observed that increased *HDAC1* expression in patient tissues is associated with similarly altered gene expression. We confirmed that *HDAC1* overexpression correlates with resistance to multiple drugs in an additional PDAC cell line (PANC-1). Using ChIP-seq, we identified regulatory sequences and nearby genes directly impacted by *HDAC1* overexpression. These genes were enriched for GTPases and the EMT pathway. Supporting the importance of these pathways in patients, we show that the expression of these genes in patient tissues was negatively correlated with overall patient survival. We used a biochemical approach to show that *HDAC1* overexpression increased GTPase activity suggesting that altered GTPase activity contributes to chemoresistance and that GTPases represent possible targets to reverse resistance. The integration of multiple genomic data types yielded insight into the role chromatin remodeling, driven by *HDAC1* overexpression, plays in drug resistance.

Using genomic analyses of an *in vitro* system perturbing HDAC1 function, we identify genes and pathways which are regulated by HDAC1 and contribute to tumorigenesis and chemoresistance. We highlight the clinical relevance of this approach by using the data to nominate a novel nine gene signature that predicts *HDAC1* expression and is associated with patient survival. We generated and analyzed multiple genomic datasets to prioritize the direct targets of HDAC1 and nominate alternative gene targets for drug development and potential markers of treatment response in patients with elevated *HDAC1* expression.

## RESULTS

### *HDAC1* overexpression induces expression of markers of EMT *in vitro* and in human PDAC tissues

We performed RNA-sequencing to measure gene expression in cell lines with *HDAC1* overexpression. We used CRISPRa^21^ to generate a stable MIA PaCa-2 cell line (MP2_HDAC1_OE) expressing *HDAC1* at approximately 3 times the levels of the control line (MP2_NTC) which expresses a non-targeting control guide (**Supplementary Fig. 1a**). MIA PaCa-2 is a well-characterized line with moderate to high expression of *HDAC1^22^. HDAC1* is the most abundantly expressed HDAC gene in this line (**Supplementary Fig. 1b**). Comparing transcriptomic profiles of MP2_HDAC1_OE and MP2_NTC cells, we found 1,259 genes that are differentially expressed with overexpression of *HDAC1* (padj < 0.1). These differentially expressed genes (DEG) are enriched for pathways involved in drug resistance: apoptosis, EMT, G2-M checkpoint, and hypoxia (**Fig. 1a, Supplementary Fig. 2**). Alteration of EMT-associated DEG promotes invasion and migration associated with a more drug resistant mesenchymal cell state^23^. The cell surface marker CD44 is characteristic of the mesenchymal phenotype^24,25^. We detected a 1.8-fold increase in *CD44* expression upon overexpression of *HDAC1*. Consistent with the expression data, immunohistochemistry showed a comparable 2-fold increase in relative density of CD44 protein in PDAC cell lines with *HDAC1* overexpression (**Fig. 1b, c**). These changes may be driven by the well-described EMT regulator ZEB1 which has increased expression in the presence of high *HDAC1* expression (L2FC=0.54, padj=0.10). No significant difference in expression was observed for *SNAI1/2* or *TWIST1* (**Supplementary Table 1**).

**Fig 1.**
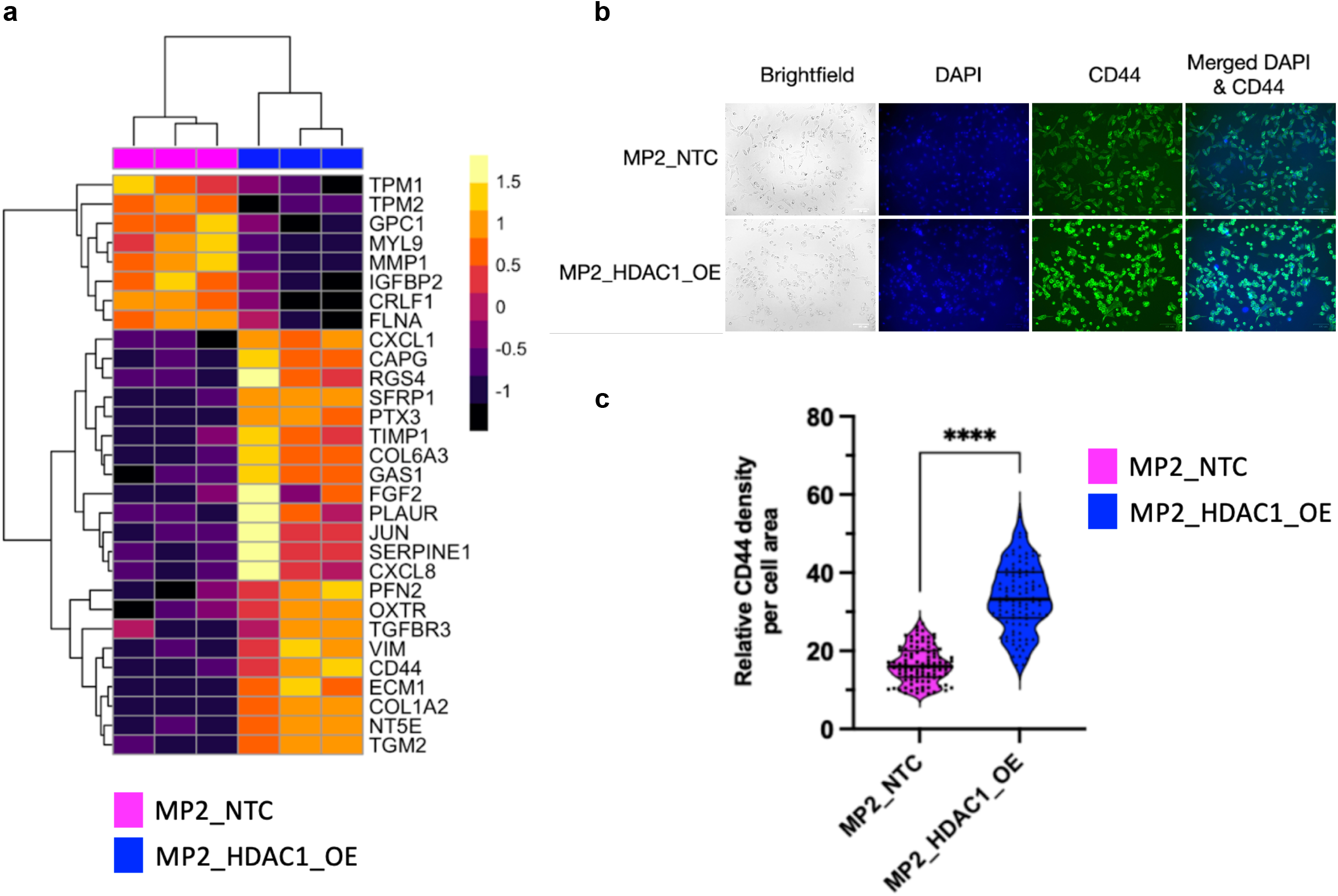
*HDAC1* overexpression is associated with increased expression of EMT genes. **a)** Expression of EMT genes in MP2_HDAC1_OE and MP2_NTC cell lines. Each column represents a replicate of the denoted cell line. The color scale denotes the z-score of each gene. **b)** Brightfield images and immunofluorescent staining of DAPI (blue), CD44 (green), and merged CD44:DAPI (blue/green) of MP2_HDAC1_OE (bottom) and MP2_NTC (top) cells. **c)** Violin plot analysis of immunofluorescent staining of CD44 in MP2_HDAC1_OE (blue) and MP2_NTC cells (pink). 100 cells were measured for each cell line. P-values were calculated using an unpaired parametric t-test. ****p<0.0001.

These data provide evidence that *HDAC1* overexpression modulates several known resistance pathways in an *in vitro* model. Next, we compared the transcriptomic effects of *HDAC1* overexpression measured *in vitro* to those observed in patient tumors. We used RNA-sequencing data collected from pancreatic tumor tissues by The Cancer Genome Atlas (TCGA-PDAC dataset). We compared gene expression from tissues with the lowest (n=45, HDAC1^LOW^) and highest (n=45, HDAC1^HIGH^) quartiles of *HDAC1* expression to identify DEG. We identified 10,592 DEG between HDAC1^HIGH^ and HDAC1^LOW^ tissues. We intersected this gene list with the 1,259 genes identified in our *in vitro* experiment and identified 322 genes that are significantly altered (padj < 0.1) in the same direction as we observed in cell lines. Heatmaps of the DEG in both datasets were clustered by sample which separated high and low *HDAC1* expression (**Fig. 2a, b**). Gene set enrichment analysis of the 322 DEG revealed multiple cancer processes including cadherin binding, cell-cell adhesion, regulation of cell migration, and GTPase activity (**Fig. 2c**, **Supplementary Table 2**). These genes represent a confident and consistent set regulated by HDAC1 *in vitro* and *in vivo*.

**Fig 2.**
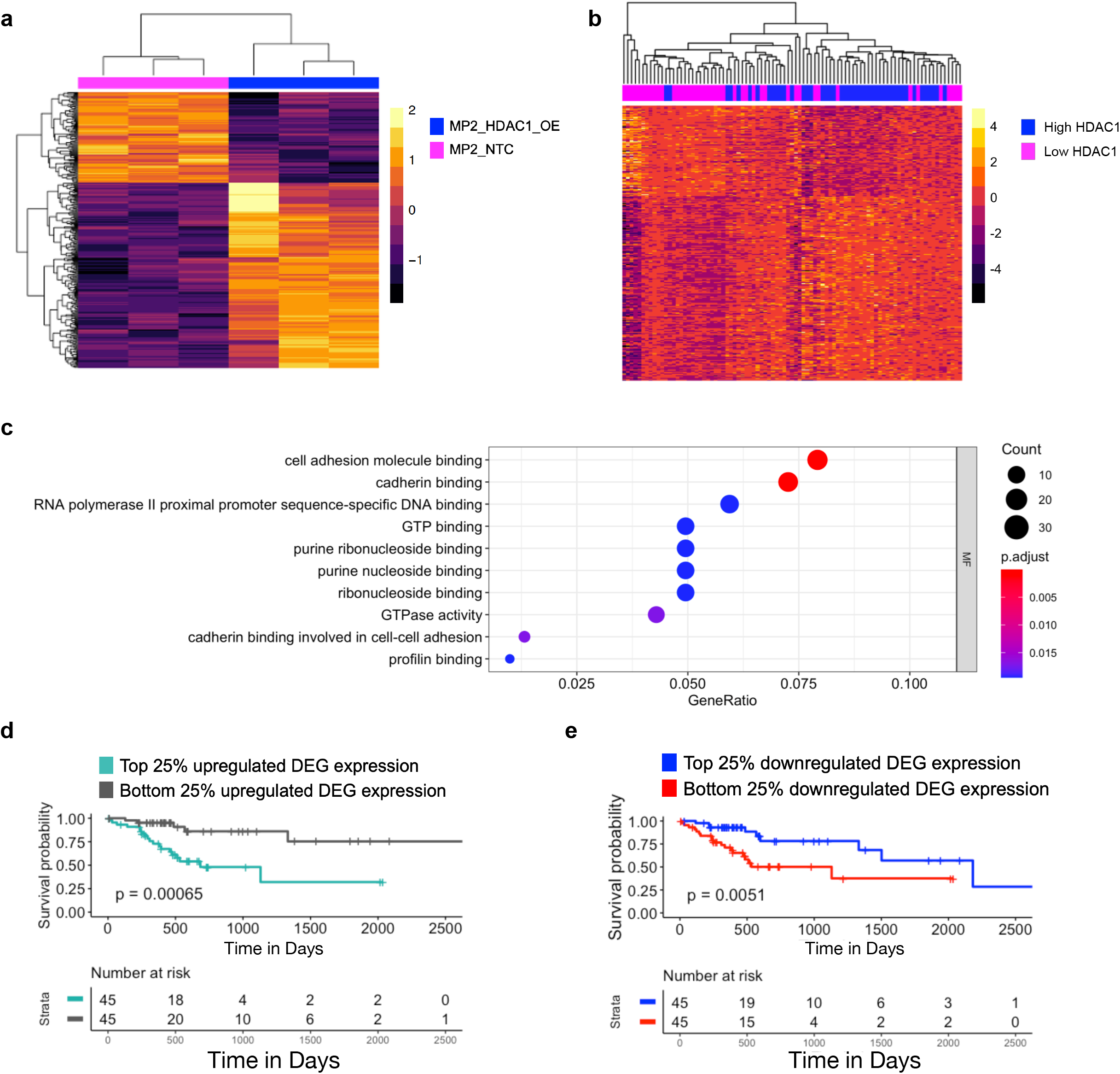
Genes altered by increased *HDAC1* expression in PDAC cell lines and TCGA PDAC samples are associated with patient survival. **a),** Expression of DEG in MP2_HDAC1_OE (blue) and MP2_NTC (pink) cell lines. DEG are significantly (padj < 0.1) altered in the same direction in MP2_HDAC1_OE cells and TCGA PDAC tissues with the top 25% of *HDAC1* expression. Each column represents a replicate of the noted cell line. The color scale denotes the z-score of each gene. **b)** Expression of DEG in TCGA PDAC samples. Each column represents a tumor sample. The color scale denotes the z-score of each gene. DEG are significantly (padj < 0.1) altered in the same direction in TCGA PDAC tissues with the top 25% of *HDAC1* expression and MP2_HDAC1_OE cells. **c)** GO analysis showing enriched molecular functions using the genes (n = 322) in **a** and **b.** **d)** Overall survival of TCCA PDAC patients (n = 90) with top and bottom 25% of average gene expression of upregulated genes (n = 216) in **a** and **b**. P-values were derived using log-rank test. **e)** Overall survival of TCCA PDAC patients (n = 90) with top and bottom 25% of average gene expression of downregulated genes (n = 106) in **a** and **b**. P-values were derived using log-rank test.

Given these data and our previous findings that linked *HDAC1* expression to cellular resistance to cytotoxic chemotherapeutic drugs^15^ we wondered whether *HDAC1* expression is associated with patient response to treatment. In the TCGA dataset, we used patient prognosis information, specifically overall survival, as a proxy for treatment response. We hypothesized that given HDAC1’s role in cellular resistance, genes associated with *HDAC1* overexpression might have prognostic value. We began with the confident set of 322 genes associated with increased *HDAC1* expression in both patient tissues and in the MP2_HDAC1_OE line. We divided these genes into two groups: upregulated genes (n = 216) and downregulated (n = 106) genes. For each gene set, we calculated a mean gene expression value from TCGA PDAC patient tissues and compared the top and bottom quartiles in a survival analysis. We determined that overall survival was shortened for patients with high expression of the upregulated genes and low expression of the downregulated genes (**Fig. 2d, e**). Although overall survival depends on multiple factors, including treatment response, this finding is consistent with our observation that *HDAC1* overexpression is associated with drug resistance *in vitro* and supports the hypothesis that these genes might also impact drug response in patients and lead to decreased survival time. Importantly, overexpression of *HDAC1* alone is not predictive of patient survival (p=0.44**, Supplementary Fig. 3a**). In an independent cohort of 26 patients^26^, we observed a similar difference in survival based on HDAC1-regulated genes (**Supplementary Fig. 3b, c**).

We tested whether our approach of combining the data we generated from cell lines with patient tumor data improved survival predictions. We compared prognostic predictions from the 322 DEG associated with *HDAC1* expression in both TCGA PDAC tumors and HDAC1_OE cell lines in **Figure 2** with the top 322 DEG in TCGA PDAC samples with high and low *HDAC1* expression as well as the top 322 DEG in our PDAC cell lines with *HDAC1* overexpression and controls. We observed a more significant p-value for overall survival when combining our *in vitro* data with patient data than when using DEG from cell lines or TCGA PDAC tumors individually (**Supplementary Fig. 4a-d**) highlighting the benefits of combining these two datasets.

### *HDAC1* overexpression leads to multi-drug resistance

We further evaluated the effects of *HDAC1* overexpression on chemoresistance by comparing drug response in MP2_HDAC1_OE and MP2_HDAC1_NTC cell lines. We also assessed the impact of *HDAC1* knockdown by treating the MIA PaCa-2 cell line with a HDAC1 DsiRNA (MP2_HDAC1_KD) (**Supplementary Fig. 5**). Under these three conditions, we measured the effect of treatment with irinotecan, gemcitabine, and oxaliplatin on cell viability (**Fig. 3a-c, Supplementary Fig. 6a-c**). MP2_HDAC1_OE cells were more resistant to drug treatment than control cells and MP2_HDAC1_KD cells. Since *HDAC1* overexpression led to increased resistance to multiple drugs, we evaluated the effect of HDAC1 protein inhibition on drug response. We treated MP2_HDAC1_OE and MP2_NTC lines with romidepsin, a HDAC1/2 inhibitor, in combination with increasing concentrations of irinotecan, gemcitabine, and oxaliplatin. We observed a sensitizing effect of romidepsin on the MP2_HDAC1_OE cells treated with each chemotherapeutic independently (**Fig. 3d, Supplementary Fig. 6d-i**). We replicated this experiment in another PDAC cell line, PANC-1, and observed the same sensitizing effect in PANC1_HDAC1_OE cells (**Supplementary Fig. 6e, g, i**). Together these experiments show a sensitizing effect of decreasing HDAC1 activity through either chemical inhibition or decreased expression.

**Fig 3.**
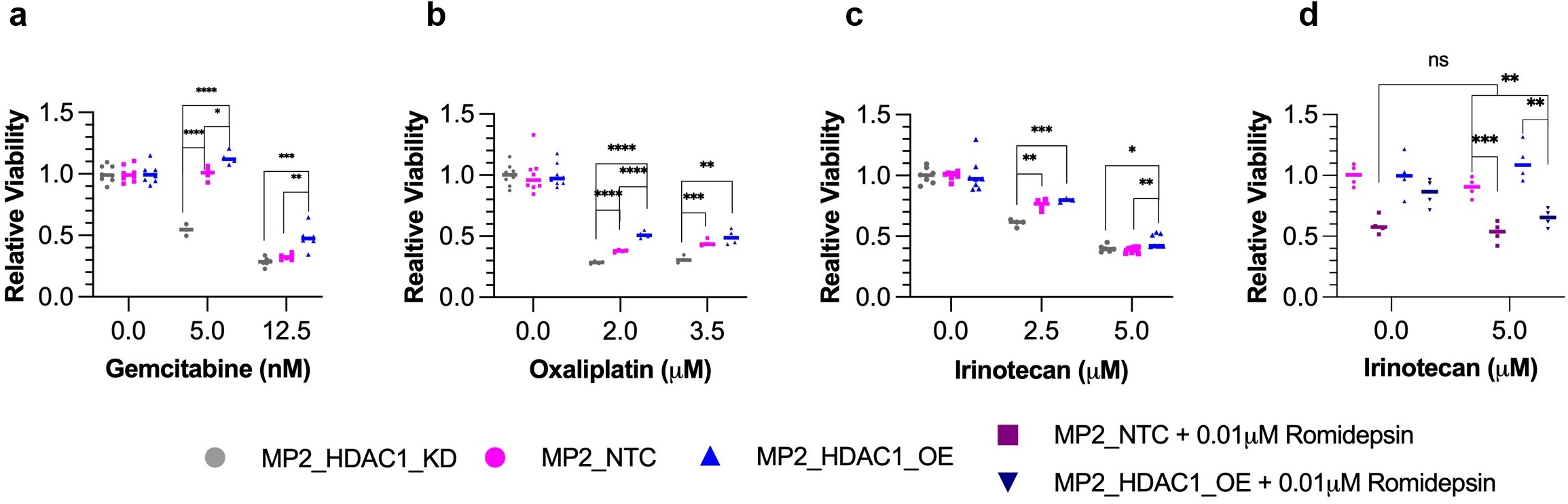
Quantification of cell viability in PDAC cell lines following treatment of chemotherapeutics. Quantification of viability following treatment with **a)** gemcitabine, **b)** oxaliplatin, and **c)** irinotecan in MP2_HDAC1_OE (blue triangles), MP2_NTC (pink squares), and MP2_HDAC1_KD (grey circles) cell lines. The bar represents the median. P-values were calculated using an unpaired parametric t-test. *p< 0.05, **p<0.01, ***p<0.001. ns, not significant. **d)** Quantification of cell viability following treatment with romidepsin, a HDAC1 inhibitor, in MP2_HDAC1_OE and MP2_NTC cell lines. Bar = median. P-values were calculated using an unpaired parametric t-test. *p< 0.05, **p<0.01, ***p<0.001. ns, not significant.

### *HDAC1* overexpression alters chromatin accessibility in distal enhancer and promoter regions nearby molecular switches

Understanding that HDAC1 inhibition has not been an effective strategy in patients, we sought to identify pathways downstream of *HDAC1* overexpression that could represent alternative targets. To identify direct and indirect impacts of *HDAC1* overexpression that might contribute to resistance, we measured genome-wide DNA binding of the HDAC1 protein and the presence of the activating histone mark, H3K27 acetylation (H3K27ac) using ChIP-sequencing in the MP2_HDAC1_OE and MP2_NTC cell lines. Using the standard ENCODE ChIP-seq protocol for peak calling^27^, we identified 17,457 binding sites for HDAC1 (10,033 unique to MP2_HDAC1_OE, 3,789 unique to MP2_NTC). We found 30,961 regions of H3K27ac; 8,392 were unique to MP2_HDAC1_OE and 5,916 were unique to MP2_NTC (**Fig. 4a**). All peaks were annotated to genomic features (i.e., promoter, distal intergenic, 5’ UTR) (**Supplementary Fig. 7**). As expected, the majority of HDAC1 binding and regions of H3K27ac occur near the transcription start sites (TSS) of DEG when *HDAC1* is overexpressed. The H3K27ac peaks specific to *HDAC1* overexpressing cells occurred significantly more near the upregulated genes despite HDAC1’s canonical role as a repressor (**Supplementary Fig. 8**).

**Fig 4.**
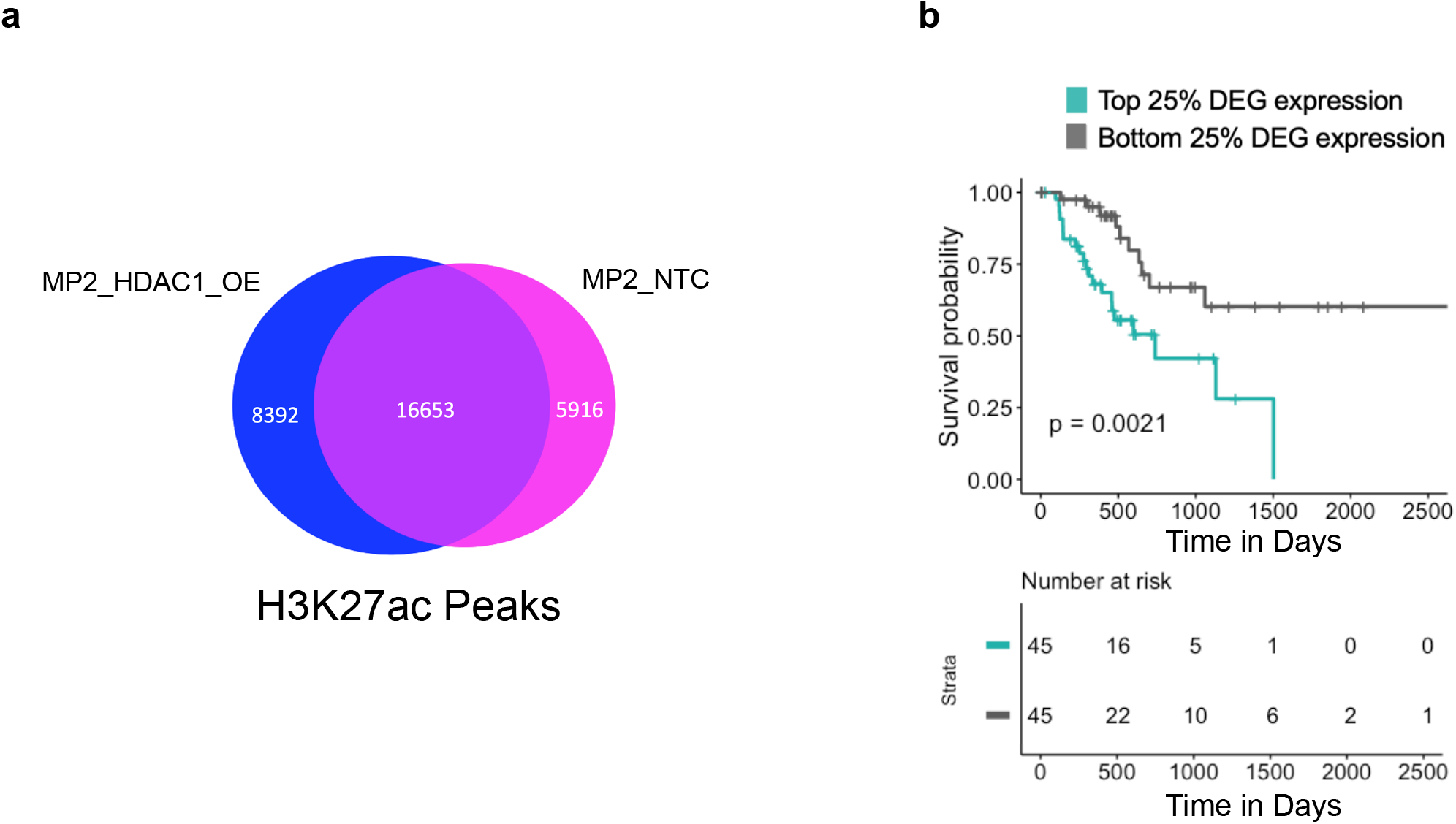
ChIP-sequencing reveals DEG with increased HDAC1 binding and H3K27 acetylation in promoter upon *HDAC1* overexpression are associated with worse patient survival. **a)** Venn diagram showing overlap of H3K27ac ChIP-seq peaks in MP2_HDAC1_OE and MP2_NTC cell lines. **b)** Overall survival of TCCA PDAC patients (n = 90) with top (teal) and bottom (grey) 25% of average gene expression of upregulated DEG with increased HDAC1 binding and H3K27 acetylation in their promoter upon *HDAC1* overexpression. P-value was derived using log-rank test.

To identify regions with altered HDAC1 binding or H3K27 acetylation directly impacting gene expression, we overlapped 1 kilobase (kb) regions centered on all HDAC1 and H3K27ac peaks (overlapping peaks were merged, see Methods) with promoter regions of DEG (2kb upstream of annotated TSS). This revealed 1,857 regions of HDAC1 binding or H3K27 acetylation in promoters of 1,040 DEG (one promoter can have more than one overlapping peak). Gene set enrichment analysis of these 1,040 DEG with evidence of direct regulation by HDAC1 revealed enrichment for GTPase activity, cadherin binding, and DNA binding (FDR < 0.05) (**Supplementary Table 3**).

To better understand the role HDAC1 plays in the regulation of PDAC pathways and especially chemoresistance, we measured how the overexpression of *HDAC1*, a histone deacetylase, impacts H3K27ac and influences gene expression. We categorized regions of HDAC1 binding and H3K27 acetylation based on whether they were increasing or decreasing across the regions described above (1kb windows centered on the peak). Using the sequencing reads collected in ChIP-seq peaks for either H3K27ac or HDAC1, we calculated a fold-change to determine whether there was evidence of increased or decreased binding with *HDAC1* overexpression. Given HDAC1’s canonical role as a repressor, we expected that increased HDAC1 binding would be associated with decreased H3K27 acetylation, however, we only identified 235 DEG with increased HDAC1 binding and reduced H3K27 acetylation in the promoter regions (+/- 2kb from TSS). In contrast, the promoters of 597 DEG had increased HDAC1 binding and increased H3K27 acetylation (fold-change > 1) upon *HDAC1* overexpression and 407 were upregulated (**Supplementary Table 4**). While this finding does go against the canonical understanding of HDAC1 as a repressor, previous studies have shown that HDAC1 binding can be found near actively transcribed genes^16^. Continuing under the assumption that these genes directly bound by HDAC1 with altered H3K27 acetylation represent an important subset of directly regulated genes, we tested whether genes whose promoters had altered HDAC1 binding and H3K27 acetylation were associated with overall patient survival. We performed survival analysis comparing outcomes of patients with the top 25% and bottom 25% mean tumor gene expression of these 597 genes. Patients with the highest mean expression of upregulated genes and lowest mean expression of downregulated genes have significantly worse overall survival (**Fig. 4b**, **Supplementary Fig. 9a**). We also showed that *HDAC1* expression is significantly higher in patients with the top 25% of mean tumor gene expression of the 597 genes (**Supplementary Fig. 9b**).

### *HDAC1* overexpression leads to increased GTPase activity

The identification of DEG upon *HDAC1* overexpression led us to explore one pathway that has not been previously linked to chemoresistance. Pathway enrichment analysis of DEG with increased HDAC1 binding and H3K27 acetylation in promoters (n = 597) identified an enrichment for many known cancer pathways (**Supplementary Table 5**). Included on this list was Ras signaling, regulation of apoptotic signaling pathway, chromatin binding, and GTPase activity. GTPase activity was also significant in enrichment analyses described in **Figure 2** driven by overexpression of the GTPases and associated proteins (e.g. *RALB, RAB27B*, and *RAC1*) which are upregulated with *HDAC1* overexpression and are associated with significantly worse overall patient survival (**Fig. 5a, b**). *RAP2B*, a ras-related GTP-binding protein, and *ARHGAP5*, a Rho family-GTPase activating protein, are examples of genes upregulated upon *HDAC1* overexpression. When activated, many of these GTP-binding proteins promote cell migration, cell adhesion, proliferation, and metastasis in cancer^28–30^ (**Supplementary Fig. 10a, b**) (**Supplementary Fig. 11a, b**). We also observed increased HDAC1 binding and H3K27 acetylation near their TSS (**Supplementary Fig. 10c, 11c**).

**Fig 5.**
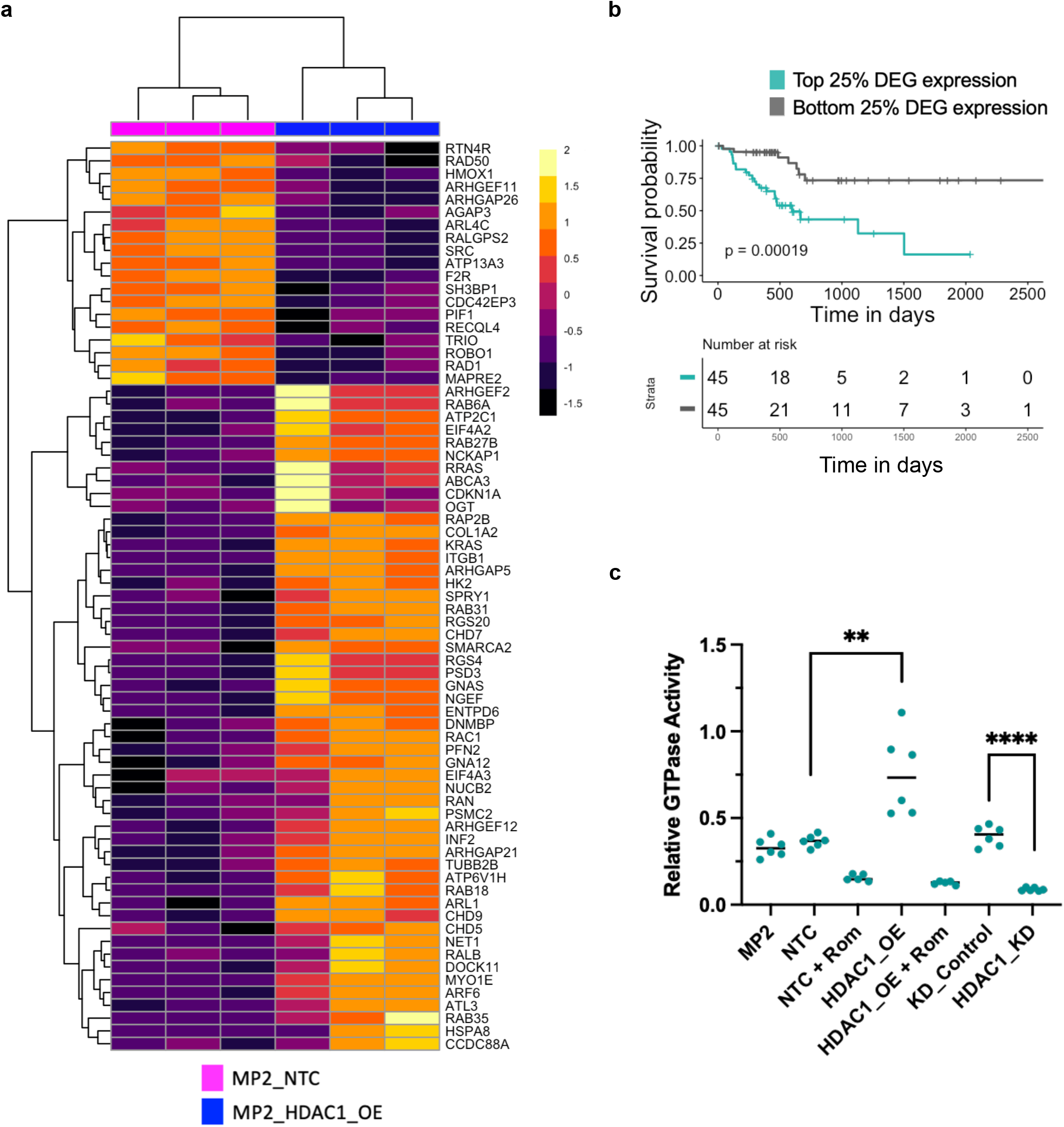
*HDAC1* overexpression is associated with increased GTPase activity. **a)** Normalized expression of genes from GO Terms that are associated with GTPases in MP2_HDAC1_OE (blue) and MP2_NTC (pink) cell lines. Each column represents a replicate of the denoted cell line. Color scale denotes z-score for each gene. **b)** Overall survival of TCCA PDAC patients (n = 90) with top (teal) and bottom (grey) 25% of average gene expression of DEG enriched for GTPase activity (genes in Supplementary Fig. 10d) when *HDAC1* is OE. P-value was derived using log-rank test. **c)** Comparison of GTPase activity in the following MIA PaCa-2 cell lines: plain MP2, NTC, NTC treated + Rom, HDAC1_OE, HDAC1_OE + Rom, KD control, and HDAC1_KD. P-values were calculated using an unpaired parametric t-test. **p< 0.01, ****p<0.0001.

While activation of KRAS is a hallmark of PDAC, less is understood about other RAS proteins identified by our analysis. Our data suggest that these proteins may also play a role in chemoresistance. We used RNA-sequencing data from 14 unmodified PDAC cells lines with varying response to gemcitabine^26^ to further support the hypothesis that GTPase activity alters cellular response to chemotherapy. We found that cell lines with increased expression of genes influencing GTPase activity (GO:0003924) had higher levels of resistance to gemcitabine (**Supplementary Fig. 11d**). We identified a total of 71 differentially expressed genes that modulate GTPase activity in the MP2_HDAC1_OE line (**Fig. 5a**), 38 of these are overexpressed in cell lines with increased resistance with 3 meeting a significant cutoff of padj < 0.05. (**Supplementary Fig. 12, Supplementary Table 6**). Increased GTPase activity activates the MAPK and PI3K pathways which promote tumor proliferation and drug resistance^31^. KEGG^32^ pathway mapping of these 597 DEG confirmed the enrichment of 21 upregulated genes in the MAPK and PI3K pathways upon *HDAC1* overexpression (**Supplementary Fig. 13a, b**).

Given the increased transcript levels of several GTPases, we tested whether there was a measurable difference in GTPase activity upon *HDAC1* overexpression. GTPase activity was measured through the detection of GTP remaining after a GTP hydrolysis reaction catalyzed by cell lysates from the MP2_HDAC1_OE line compared to the MP2_NTC line. MP2_HDAC1_OE cell lysates have significantly increased GTPase activity compared to the MP2_NTC control line. Conversely, treatment of MP2_HDAC1_OE and MP2_NTC cells with romidepsin, a HDAC1/2 inhibitor, decreased GTPase activity. We also observed decreased GTPase activity in MIA PaCa-2 cells with a DsiRNA targeting HDAC1 (MP2_HDAC1_KD) compared to MIA PaCa-2 cells with a non-targeting DsiRNA (KD_Control) (**Fig. 5c, Supplementary Table 7**). These data demonstrate that *HDAC1* overexpression increases GTPase activity and that inhibition of HDAC1 reverses the effect.

### Expression of 9 HDAC1-regulated genes predicts PDAC patient survival

Identification of prognostic signatures in PDAC could be of clinical utility because tumor classification can improve guidance for therapeutic decision making and developing a personalized treatment plan. The above analysis overlapping expression and ChIP-seq data identified 597 genes regulated by HDAC1 that are associated with patient outcomes using genes identified from *in vitro* and *in vivo* signatures of *HDAC1* overexpression, although expression of *HDAC1* alone is not prognostic. We calculated a simplified signature of patient prognosis using a multivariate logistic regression with L1 penalized log partial likelihood (LASSO) for feature selection^33^. From the 597 genes, LASSO identified a 9-transcript model sufficient to differentiate TCGA PDAC tumors with high and low *HDAC1* expression (**Fig. 6a, b, Supplementary Table 8**). To determine the clinical relevance of the genes selected using the LASSO model, survival analysis was performed comparing the patients in the top and bottom quartile of predictor values from the regression and the patients group with the highest predictor values had significantly worse overall survival (**Fig. 6c**). In order to evaluate the performance of the LASSO model we generated the area under the ROC curve (AUC) and found that the validation cohort had an AUC of 0.97 indicating that it performed as an excellent predictor model for *HDAC1* expression in patients (**Fig. 6a, b**). Even though *HDAC1* expression alone is not prognostic of survival, we were able to use the predictor values generated by this model to show that patients predicted to have higher *HDAC1* expression had significantly worse overall survival.

**Fig 6.**
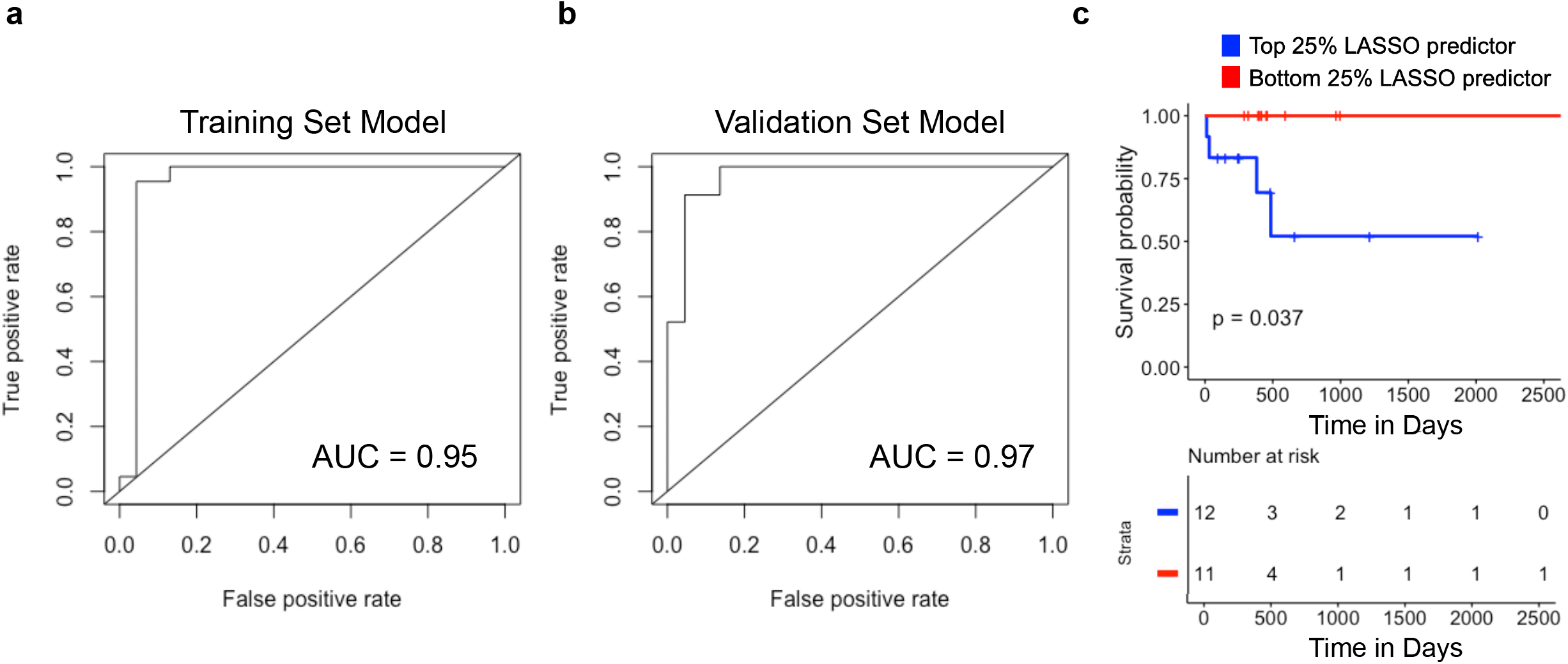
ChIP-sequencing reveals DEG associated with predictive gene signature for patient survival with high *HDAC1* expression. **a,b)** Receiver operating characteristic (ROC) curve for the nine-transcript LASSO model. Model performance on the **a)** training cohort and **b)** validation cohort. **c)** Overall survival of TCCA PDAC patients (n = 90) with top (blue) and bottom (red) 25% of predictor values generated from the nine transcript LASSO model. P-value was derived using log-rank test.

## DISCUSSION

Pancreatic cancer ranks among the deadliest cancers due to its chemoresistant nature and insufficient treatment options. Understanding what drives chemoresistance is essential to identifying new therapeutic targets and improving patient outcomes. Chromatin remodeling has been established as a critical feature of tumorigenesis and cancer progression, making the pathway an attractive drug target. Using genomic and biochemical approaches we revealed potential mechanisms by which *HDAC1* overexpression contributes to chemoresistance and showed that HDAC1 inhibition sensitizes PDAC cells to chemotherapeutic treatment, further strengthening the argument that this pathway is a good candidate for targeted treatment, however HDAC inhibitors have faced challenges in clinical trials. Commercial HDAC inhibitors target a class of HDACs rather than specific proteins. This cross-reactivity leads to genome-wide off target effects and patient toxicity. That has motivated the current study which aims to better understand how *HDAC1* activation contributes to resistance and reveal novel downstream targets that may lead to alternative treatment strategies. The integration of multiple genomic datasets has enabled the successful nomination of at least one novel therapeutic approach.

In contrast to other tumor types, multiple large-scale drug trials that used targeted therapy were not as successful in pancreatic cancer^34,35^, thus using a targeted gene panel that can be used to better define potential treatment options for PDAC patients could lead to improved survival and quality of life^36,37^. In this study, we collected data from an *in vitro* system testing the impact of *HDAC1* overexpression on PDAC cells and combined these results with information from publicly available gene expression data gathered from both tissues of PDAC patients and PDAC cell lines to show that *HDAC1* overexpression regulates a set of transcriptomic responses that contribute to chemoresistance and a signature of genes regulated by HDAC1 can also be shown to predict *HDAC1* expression and is associated with patient outcome. *HDAC1* overexpression alone is not significantly prognostic of worse overall survival in PDAC patients, however, the genes altered by *HDAC1* overexpression are prognostic. This suggests that only a subset of HDAC1-regulated pathways affect outcomes. Our results explore the pathways under the control of HDAC1 that contribute to patient survival and show that they can be used to predict outcomes that may be linked to treatment response.

We integrated several datasets to better understand how *HDAC1* overexpression impacts PDAC cells. Beginning with transcriptomic data, we clearly show that *HDAC1* overexpression impacts several processes, including EMT, known for their role in tumorigenesis, progression, and drug resistance. Cells that undergo EMT also have a more stem cell-like phenotype and are associated with suppression of proteins involved in drug transport, such as CNT3, allowing the cells to evade the anti-proliferative effects of chemotherapeutics including gemcitabine^5,38^. Increased expression of *CD44*, a cell surface protein important for cell adhesion and migration, is associated with a more mesenchymal-like phenotype which is characteristic of EMT^24^. Here we have shown that the mesenchymal marker, CD44 transcript and protein are more abundant in cells with *HDAC1* overexpression, agreeing with our past work showing that *HDAC1* overexpression leads to increased migration^15^.

Induction of EMT is also associated with drug resistance and we showed that cells overexpressing *HDAC1* are resistant to multiple drugs. Characterizing the direct regulatory impacts of HDAC1 binding and H3K27ac occupancy allowed us to prioritize the genes most likely involved in patient outcomes. Our ChIP-seq experiments revealed altered H3K27 acetylation and HDAC1 binding nearby genes whose expression changed with *HDAC1* overexpression. Interestingly, we found that the majority of the DEG with HDAC1 binding had an increase in HDAC1 and H3K27ac signals near their promoter. Our findings are in agreement with a previously published study concluding that HDAC1 binding is enriched at actively transcribed genes^16^. It is still not well understood how HDAC1 binding activates gene expression but others have shown that HDAC1 binding can regulate RNA Pol II recruitment at promoters and that HDAC1 may regulate transcription elongation through interaction with BRD4^17^. Additionally, HDAC1 is known to deacetylate proteins other than histones which might facilitate activation of nearby transcription factors (e.g. MYC)^17^. HDACs are enriched distally at super enhancers, in addition to promoters, but their function remains poorly understood outside of their role in repression^20^. While our data showed the majority of HDAC1 binding in promoter regions, a better understanding of HDACs role in enhancer regions is also important for a full understanding of the impact of HDAC1 dysregulation.

Throughout this study, our gene set enrichment analyses of genes associated with *HDAC1* overexpression consistently revealed GTPase activity as an enriched process. The well-described driver of PDAC, KRAS is activated in almost all tumors (including the MP2 and Panc1 cell lines), however these data suggest other Ras proteins and their regulators and partners also play a role. A variety of proteins contribute to GTPase activity and many are druggable^39^, which makes them of potential clinical interest. We have shown that expression of GTPases and GTPase activating proteins are associated with significantly worse overall survival. We used biochemical assays to confirm that cells overexpressing *HDAC1* have higher GTPase activity than control cells. This effect was reversible in cells treated with a HDACi, which reduced GTPase activity. GTPase signaling is important for pancreatic cancer initiation, metastasis, and invasion^40^. Increased GTPase signaling leads to the activation of key signaling cascades, such as MAPK and PI3K, that regulate cell proliferation, migration, and drug resistance in cancer^31^. We have shown that the expression of genes in the MAPK and PI3K pathways are increased upon *HDAC1* overexpression, highlighting known pathways altered by HDAC1 that contribute to PDAC progression^41^. GTPases, such as RAC1, can activate EMT in multiple cancers, thus leading to a more invasive and drug resistant phenotype^42,43^ and we show that *RAC1* expression is increased upon *HDAC1* overexpression in PDAC cells. Expression of *ARHGAP5*, a GTPase activating protein which also promotes EMT^30^, and of several other proteins promoting GTPase activity are increased upon *HDAC1* overexpression and their transcripts are more abundant in gemcitabine resistant PDAC cell lines.

Finally, we narrow the list of genes impacted by *HDAC1* overexpression to a novel 9-transcript signature associated with *HDAC1* expression that successfully predicts patient survival. This highlights the potential clinical utility of data generated *in vitro* in predicting and understanding molecular mechanisms of disease in patients. Our panel of potential biomarkers represents a step forward in the development of an assay that is predictive of patient survival and which could influence treatment decisions.

Despite decades of research, the treatment of PDAC patients relies largely on cytotoxic chemotherapies which have limited effectiveness in treating late-stage disease. Identifying patients who will benefit from existing treatments or those who need an alternative treatment is a key clinical need. Our genomic analyses identified a role for HDAC1 in regulation of transcriptional programs that are relevant for patient outcomes and have nominated novel therapeutic strategies for individuals who are predicted to experience poor outcomes and chemotherapeutic resistance. This knowledge will be key as the field of oncology continues to implement precision medicine.

## MATERIALS AND METHODS

### Cell Culture

MIA PaCa-2 cells (CVCL_0428, ATCC #CRM-CRL-1420) and PANC-1 cells (CVCL_0480, ATCC #CRM-CRL-1469) were cultured in D10 media: DMEM (Lonza #12-614Q) supplemented with 10% FBS (GELifeSciences #SH30071.03), and 0.5% penicillin-streptomycin (ThermoFisher #15140122). All cell lines were maintained at 37 °C and 5% CO_2_. Cells were cryopreserved with the addition of 10% DMSO (EMD #MX1458-6).

### Plasmids

LentiCRISPRv2 (Addgene #52961) or lentiSAMv2 (Addgene #92062) and lenti-MS2-p65-HSF1-Hygro (Addgene #89308) were used to generate stable cell lines for gene knockout and activation, respectively. pMD2.G (Addgene #12259) and psPAX2 (Addgene #12260) were used to facilitate viral packaging of sgRNAs and single vector plasmids.

### sgRNA Cloning

gRNA oligos were designed and cloned into their respective plasmids as described previously^21^.

### DsiRNA

IDT TriFECTa RNAi kit was used per manufacturer’s protocol. 100,000 cells were seeded in 1 well of a 12 well tissue culture treated plate 24 hours prior to transfection. Cells were transfected using RNAiMax (ThermoFisher #13778-030) following manufacturer’s recommended protocol. As indicated in the TriFecta kit (IDT hs.Ri.HDAC1.13.2-SEQ1: 5’-rCrUrGrGrArArCrUrGrCrUrArArArGrUrArUrCrArCrCrAGA-3’, hs.Ri.HDAC1.13.2-SEQ2: 5’-rUrCrUrGrGrUrGrArUrArCrUrUrUrArGrCrArGrUrUrCrCrArGrGrA-3’), TYE 563 transfection efficiency control, positive HPRT-S1 control, and negative (IDT DS NC1) scrambled sequence control were utilized. Further assays were performed 48 hours after transfection. Expression was validated with each transfection with the IDT PrimeTime qPCR Assay system (HDAC1 Exon 1-2 Hs.PT.58.20534173, HDAC1 Exon 3-4 Hs.PT.58.38680914, ACTB Exon 1-2 Hs.PT.39a.22214847, GAPDH Exon 2-3 Hs.PT.39a.22214836, HPRT1 Exon 8-9 Hs.PT.58v.45621572) on an Agilent QuantStudio 6 Flex Real-Time PCR system.

### GTPase-Glo Assay Using Cell Lysates

*In vitro* GTPase activity was measured using the GTPase-Glo assay (Promega #V7681). We followed the protocol as described^44^ with modifications for use of cell lysates. Cell lysates were made from the following MIA PaCa-2 cell lines: MP2, MP2_HDAC1_OE, MP2_NTC, MP2 with DsiRNA targeting *HDAC1* (HDAC1_KD), MP2 with a non-targeting DsiRNA (KD_Control), MP2_HDAC1_OE treated with 0.01 μM romidepsin, an HDAC1 inhibitor, and MP2_NTC treated with 0.01 μM romidepsin. Cells (2 x 10^6^ per tube) were lysed in a lysis buffer containing 50 mM HEPES at pH 7.6, 150 mM NaCl, 10% Glycerol, 0.1% NP-40, and 2 mM MgCl_2_. To generate the lysate, 10μL of lysis buffer per 100,000 cells was added to each cell pellet and resuspended. Lysates were mixed for 30 minutes at 4°C, vortexed in three 10 second intervals, then centrifuged at 4°C for 30 minutes at 16.1x RCF. A 2X GTP solution was prepared and the reaction was initiated following the manufacturer’s protocol.

Modifications for cell lysates required background wells for each cell line. GTPase-Glo Buffer was added to cell lysates at a final concentration of 1 μL per 10,000 cells. After the GTPase reaction, 20μL was added to each respective background well. Luminescence was measured using a BioTek Synergy H5 plate reader. To calculate GTPase activity for each cell type, we calculated the difference between the luminescence of the experimental wells and background wells. GraphPad Prism 9 (version 9.3.1) was used for plotting bar charts and t-tests performed in GraphPad were unpaired, parametric, two-tailed with 95% confidence interval.

### ChIP-sequencing

MP2_HDAC1_OE and MP2_NTC cells (2 x 10^7^) were cross-linked, harvested, and DNA was precipitated using a commercial H3K27ac antibody (Abcam, ab4729). Libraries were constructed, pooled, and sequenced using an Illumina NovaSeq instrument with 75bp single-end reads. These data were generated and analyzed using published ENCODE protocols^27^ (https://www.encodeproject.org/documents/).

Differential binding analysis was conducted using the “multiBigwigSummary’’ tool from the “deepTools’’ package^45^. Using this tool, a ChIP-seq score was generated for each sample and region using genomic coordinates defined as +/- 500 bp from the center of peaks defined using the published ENCODE protocol^27^ and 1 kb upstream of all annotated genes. Regions were merged together if they overlapped. We omitted any regions with a ChIP-seq score less than 1 for both MP2_HDAC1_OE and MP2_HDAC1_NTC. Using the ChIP-seq score, we calculated a fold-change between MP2_HDAC1_OE and MP2_NTC for HDAC1 and H3K27ac in each defined region. Bound regions were categorized based on a fold-change greater than or less than one for HDAC1 binding and H3K27 acetylation.

### 3’ RNA-sequencing

Cell pellets were frozen at −80°C until RNA extraction. For RNA extraction 350 μl of RL Buffer plus 1% β-ME from the Norgen Total RNA extraction kit was added to each cell pellet and extraction proceeded per manufacturer’s instructions including use of the DNase kit (Norgen # 37500, 25720). RNA quality was verified with the Agilent BioAnalyzer RNA Nano 600 kit (cat# 5067-1512) with the RIN range between 9.2-10. RNA-sequencing libraries were made using Lexogen QuantSeq 3’ mRNA-Seq Library Prep Kit FWD for Illumina kit (cat# 015.24) with 250 ng of RNA input. They were pooled and sequenced on an Illumina NextSeq 500 instrument with 75 bp single-end reads. Read counts averaged 4 million reads and an average Q30 of 91.28%. Lexogen’s BlueBee integrated QuantSeq data analysis pipeline was used for trimming, mapping, and alignment and the R package “DESeq2”^46^ was used for differential expression analysis.

### Drug resistance screening

Cells were seeded in 96-well plates at 2000 cells/well. Seeded cells were dosed with a range of concentrations of each drug: gemcitabine (0-12.5nM), oxaliplatin (0-3.5μM), or irinotecan (0-5μM).

Cells were given a second dose of drug at the same concentration as the first 48 hours later. The number of viable cells surviving drug treatment were assayed with CellTiter-Glo (Promega #G7571) 24 hours after the last drug treatment per manufacturer’s protocol using a BioTek Synergy H5 plate reader.

HDAC1 inhibition with romidepsin (Sigma #SML1175-1MG) was performed similarly to above except that cells were dosed every 24 hours with either 0.01 μM romidepsin or with a range of irinotecan, oxaliplatin, or gemcitabine. Equal volume DMSO was used as a control in place of romidepsin. The number of viable cells surviving drug treatments were assayed with CellTiter-Glo (Promega #G7571) 24 hours after the last drug treatment per manufacturer’s protocol using a BioTek Synergy H5 plate reader. In both cases, data were plotted using GraphPad Prism 9, version 9.3.1. T-tests performed were unpaired, parametric, two-tailed with 95% confidence interval.

### Cell staining

75,000 MIA PaCa-2 cells with non-targeting, HDAC1 OE sgRNAs, or HDAC1 KD with DsiRNA were seeded in 12 well plates. Cells were stained using Alexa Fluor 488 Conjugate kit for live cell imaging (LifeTechnologies #A25618) for CD44 via the manufacturer’s protocol. DAPI (Invitrogen #D21490) was counterstained per manufacturer’s protocols for adherent cells. Presence of CD44 in the cells was quantified using ImageJ 1.53K with measurements (area, mean, and integrated density) for stain and background taken with the freehand selection tool. Relative CD44 intensity or bound CD44 per area was calculated for each cell (100 cells total per type) by: integrated density of cell-integrated density of background for that cell/area of that cell. GraphPad Prism 9 (version 9.3.1) was used for plotting violin plots and t-tests performed in GraphPad were unpaired, parametric, two-tailed with 95% confidence interval.

### Enrichment Analysis

Enrichr, a comprehensive gene set analysis web server, and the R package ClusterProfiler (version 3.12.0)^47^ were used for enrichment analysis of the differentially expressed genes^48^. We focused on the pathways (MSigDB) and gene ontology molecular function and biological process terms (GO MF, GO BP) reaching the significance threshold of FDR < 0.05.

The GO terms used to select genes in **Fig. 5a** were GDP Binding (GO:0019003), GTPase activity (GO:0003924), NTPase activity (GO:0017111), regulation of small GTPase signal transduction (GO:0051056), positive regulation of ras signal transduction (GO:0046579), and small GTPase signal transduction (GO:0007264).

### Survival Analysis

To conduct survival analysis, clinical and RNA-seq expression data was retrieved from The Cancer Genome Atlas (TCGA) for 178 PDAC (TCGA-PAAD) patients (https://portal.gdc.cancer.gov/). Data was normalized using the R package DESeq2^46^ and differentially expressed genes with an FDR < 0.1 were used to generate Kaplan-Meier survival curves. We classified tissues based on their mean expression of a given gene set (bottom, middle, and top quartiles of gene expression). We compared the patients with the lowest and highest quartile of mean gene expression and performed survival analysis. Survival curves and analyses were generated using the ggplot2, survminer, and survival R packages^49–51^.

### LASSO model selection

A predictive gene signature from transcripts that are differentially expressed (DESeq2 FDR < 0.1) and have increased HDAC1 binding and H3K27 acetylation near their TSS (+/- 2000 bp) was developed using the LASSO regression model. LASSO was performed using the R package glmnet (version 4.1-3)^52^. The TCGA PDAC cohort was split into three groups by *HDAC1* expression (top 25%, middle 50%, and bottom 25%). The cohort was further subset by randomly distributing an equal number of samples from the top 25% and bottom 25% of *HDAC1* expression into two groups (n = 45). The training cohort and the validation cohort used the same dichotomization threshold (top 25% and bottom 25% of *HDAC1* expression). Model performance was evaluated based on the model’s ability to classify patients into the high or low *HDAC1* expression group. We generated an area under the curve (AUC) value using the R package ROCR (version 1.0-11)^53^. Kaplan-Meier curves were generated using the R package survival (version 3.2-13)^50^.

### Annotation of Genomic Features

ChIP-sequencing IDR peaks were annotated to genomic features (i.e., promoter, distal intergenic, 5’ UTR) using the annotatePeak tool and then visualized using the plotAnnoBar tool from the R package ChIPseeker (version 1.27.2)^54^. TSS regions were defined as −2kb to +1kb.

### RNAseq Data Analysis

An adjusted p-value < 0.05 was used to identify differentially expressed genes from RNA-sequencing data. Using the R package DESeq2 (version 1.24.0)^46^, differentially expressed genes were excluded from the analysis if baseMean <10.

## Data Availability

ChIP-sequencing data is available using the GEO accessions GSE209895 (H3K27ac) and GSE158541 (HDAC1). RNA-sequencing data are available using the GEO accessions GSE79668 (PDAC tissues) and GSE79669 (gemcitabine resistant and sensitive cell lines)^26^. Clinical data and RNA-sequencing data for TCGA PDAC samples, 178 samples in this cohort that had matched clinical and RNA-sequencing data, were retrieved on 04/01/2020 using the GDC Data Portal (https://portal.gdc.cancer.gov/).

## Code Availability

All the code used for data analysis and generation of figures will be available upon request. Software used includes: GraphPad Prism 9 and R packages: survival version 1.2.1335^50^, survminer version 0.4.9^51^, ggplot2 version 3.3.6^49^, DESeq2 version 1.24.0^46^, pheatmap version 1.0.12^55^, clusterProfiler version 3.12.0^47^, glmnet version 4.1-3^52^, ROCR version 1.0-11^53^, ChIPseeker version 1.27.2^54^, deepTools version 3.5.0^45^, and IGV version 2.7.2^56^.

## Acknowledgements

We thank the Myers and Cooper labs for helpful feedback. We also acknowledge the data produced by The Cancer Genome Atlas which were extremely valuable and without which this study would not be possible. SJC is supported by UL1TR003096. SJC, ERG, and CAW are supported by Alabama’s State Cancer Fund.

## Author information

### Authors and Affiliations

The University of Alabama in Huntsville, Huntsville, AL, 35899, USA

Carter A. Wright

HudsonAlpha Institute for Biotechnology, Huntsville, AL, 35806, USA

Carter A. Wright, Emily R. Gordon, Sara J. Cooper

### Contributions

CAW, ERG, and SJC designed the experiments. CAW and ERG collected data. CAW and ERG analyzed the data. CAW and SJC wrote the first draft. All authors contributed to the writing of the paper and read and approved the final manuscript.

### Corresponding author

Correspondence to Sara J Cooper

### Ethics Declarations

The authors declare no competing interests.

## REFERENCES

1. Cancer of the pancreas - cancer stat facts. SEER https://seer.cancer.gov/statfacts/html/pancreas.html.

2. Hontecillas-Prieto, L. et al. Synergistic Enhancement of Cancer Therapy Using HDAC Inhibitors: Opportunity for Clinical Trials. Front. Genet. 11, 578011 (2020).

3. Bondarev, A. D. et al. Recent developments of HDAC inhibitors: Emerging indications and novel molecules. Br. J. Clin. Pharmacol. 87, 4577–4597 (2021).

4. Laschanzky, R. S. et al. Selective Inhibition of Histone Deacetylases 1/2/6 in Combination with Gemcitabine: A Promising Combination for Pancreatic Cancer Therapy. Cancers 11, (2019).

5. Palamaris, K., Felekouras, E. & Sakellariou, S. Epithelial to Mesenchymal Transition: Key Regulator of Pancreatic Ductal Adenocarcinoma Progression and Chemoresistance. Cancers 13, (2021).

6. Xu, Y. & Villalona-Calero, M. A. Irinotecan: mechanisms of tumor resistance and novel strategies for modulating its activity. Ann. Oncol. 13, 1841–1851 (2002).

7. Beatty, G. L., Werba, G., Lyssiotis, C. A. & Simeone, D. M. The biological underpinnings of therapeutic resistance in pancreatic cancer. Genes Dev. 35, 940–962 (2021).

8. Shah, V. M., Sheppard, B. C., Sears, R. C. & Alani, A. W. Hypoxia: Friend or Foe for drug delivery in Pancreatic Cancer. Cancer Lett. 492, 63–70 (2020).

9. O’Brien, M. A. & Kirby, R. Apoptosis: A review of pro-apoptotic and anti-apoptotic pathways and dysregulation in disease. J. Vet. Emerg. Crit. Care 18, 572–585 (2008).

10. Hasan, N. & Ahuja, N. The Emerging Roles of ATP-Dependent Chromatin Remodeling Complexes in Pancreatic Cancer. Cancers 11, (2019).

11. Ying, H. et al. Genetics and biology of pancreatic ductal adenocarcinoma. Genes Dev. 30, 355–385 (2016).

12. McCleary-Wheeler, A. L. et al. Insights into the epigenetic mechanisms controlling pancreatic carcinogenesis. Cancer Lett. 328, 212–221 (2013).

13. Li, Y. & Seto, E. HDACs and HDAC Inhibitors in Cancer Development and Therapy. Cold Spring Harb. Perspect. Med. 6, (2016).

14. Park, S.-Y. & Kim, J.-S. A short guide to histone deacetylases including recent progress on class II enzymes. Exp. Mol. Med. 52, 204–212 (2020).

15. Ramaker, R. C. et al. Pooled CRISPR screening in pancreatic cancer cells implicates co-repressor complexes as a cause of multiple drug resistance via regulation of epithelial-to-mesenchymal transition. BMC Cancer vol. 21 Preprint at https://doi.org/10.1186/s12885-021-08388-1 (2021).

16. Wang, Z. et al. Genome-wide mapping of HATs and HDACs reveals distinct functions in active and inactive genes. Cell 138, 1019–1031 (2009).

17. Greer, C. B. et al. Histone Deacetylases Positively Regulate Transcription through the Elongation Machinery. Cell Rep. 13, 1444–1455 (2015).

18. Chen, C. et al. The histone deacetylase HDAC1 activates HIF1α/VEGFA signal pathway in colorectal cancer. Gene 754, 144851 (2020).

19. Hai, R., He, L., Shu, G. & Yin, G. Characterization of Histone Deacetylase Mechanisms in Cancer Development. Front. Oncol. 11, 700947 (2021).

20. Gryder, B. E. et al. Chemical genomics reveals histone deacetylases are required for core regulatory transcription. Nat. Commun. 10, 3004 (2019).

21. Joung, J. et al. Genome-scale CRISPR-Cas9 knockout and transcriptional activation screening. Nat. Protoc. 12, 828–863 (2017).

22. Minami, F. et al. Morphofunctional analysis of human pancreatic cancer cell lines in 2-and 3-dimensional cultures. Sci. Rep. 11, 6775 (2021).

23. Bulle, A. & Lim, K.-H. Beyond just a tight fortress: contribution of stroma to epithelial-mesenchymal transition in pancreatic cancer. Signal Transduct Target Ther 5, 249 (2020).

24. Zhao, S. et al. CD44 Expression Level and Isoform Contributes to Pancreatic Cancer Cell Plasticity, Invasiveness, and Response to Therapy. Clin. Cancer Res. 22, 5592–5604 (2016).

25. Luu, T. Epithelial-Mesenchymal Transition and Its Regulation Mechanisms in Pancreatic Cancer. Front. Oncol. 11, 646399 (2021).

26. Kirby, M. K. et al. RNA sequencing of pancreatic adenocarcinoma tumors yields novel *expression patterns associated with long-term survival and reveals a role for ANGPTL4*. Mol.Oncol. 10, 1169–1182 (2016).

27. Landt, S. G. et al. ChIP-seq guidelines and practices of the ENCODE and modENCODE consortia. Genome Res. 22, 1813–1831 (2012).

28. Li, J. et al. TMEM43 promotes pancreatic cancer progression by stabilizing PRPF3 and regulating RAP2B/ERK axis. Cell. Mol. Biol. Lett. 27, 24 (2022).

29. Di, J. et al. Rap2B promotes proliferation, migration, and invasion of human breast cancer through calcium-related ERK1/2 signaling pathway. Sci. Rep. 5, 12363 (2015).

30. Tian, T. et al. Investigation of the role and mechanism of ARHGAP5-mediated colorectal cancer metastasis. Theranostics 10, 5998–6010 (2020).

31. Soriano, O., Alcón-Pérez, M., Vicente-Manzanares, M. & Castellano, E. The Crossroads between RAS and RHO Signaling Pathways in Cellular Transformation, Motility and Contraction. Genes 12, (2021).

32. Kanehisa, M., Furumichi, M., Sato, Y., Ishiguro-Watanabe, M. & Tanabe, M. KEGG: integrating viruses and cellular organisms. Nucleic Acids Res. 49, D545–D551 (2021).

33. Tibshirani, R. Regression shrinkage and selection via the lasso. J. R. Stat. Soc. 58, 267–288 (1996).

34. Hecht, J. R. et al. Randomized Phase III Study of FOLFOX Alone or With Pegilodecakin as Second-Line Therapy in Patients With Metastatic Pancreatic Cancer That Progressed After Gemcitabine (SEQUOIA). J. Clin. Oncol. 39, 1108–1118 (2021).

35. Tempero, M. et al. Ibrutinib in combination with nab-paclitaxel and gemcitabine for first-line treatment of patients with metastatic pancreatic adenocarcinoma: phase III RESOLVE study. Ann. Oncol. 32, 600–608 (2021).

36. Hosein, A. N., Dougan, S. K., Aguirre, A. J. & Maitra, A. Translational advances in pancreatic ductal adenocarcinoma therapy. Nat Cancer 3, 272–286 (2022).

37. Wei, X. et al. A 14-gene gemcitabine resistance gene signature is significantly associated with the prognosis of pancreatic cancer patients. Sci. Rep. 11, 6087 (2021).

38. Adamska, A. et al. Molecular and cellular mechanisms of chemoresistance in pancreatic cancer. Adv. Biol. Regul. 68, 77–87 (2018).

39. Prieto-Dominguez, N., Parnell, C. & Teng, Y. Drugging the Small GTPase Pathways in Cancer Treatment: Promises and Challenges. Cells 8, (2019).

40. Yoshimachi, S. et al. Ral GTPase-activating protein regulates the malignancy of pancreatic ductal adenocarcinoma. Cancer Sci. 112, 3064–3073 (2021).

41. Mehra, S., Deshpande, N. & Nagathihalli, N. Targeting PI3K Pathway in Pancreatic Ductal Adenocarcinoma: Rationale and Progress. Cancers 13, (2021).

42. Zhou, Y. et al. Rac1 overexpression is correlated with epithelial mesenchymal transition and predicts poor prognosis in non-small cell lung cancer. J. Cancer 7, 2100–2109 (2016).

43. Ungefroren, H., Witte, D. & Lehnert, H. The role of small GTPases of the Rho/Rac family in TGF-β-induced EMT and cell motility in cancer. Developmental Dynamics vol. 247 451–461 Preprint at https://doi.org/10.1002/dvdy.24505 (2018).

44. Mondal, S., Hsiao, K. & Goueli, S. A. A Homogenous Bioluminescent System for Measuring GTPase, GTPase Activating Protein, and Guanine Nucleotide Exchange Factor Activities. Assay Drug Dev. Technol. 13, 444–455 (2015).

45. Ramírez, F., Dündar, F., Diehl, S., Grüning, B. A. & Manke, T. deepTools: a flexible platform for exploring deep-sequencing data. Nucleic Acids Res. 42, W187–91 (2014).

46. Love, M. I., Huber, W. & Anders, S. Moderated estimation of fold change and dispersion for RNA-seq data with DESeq2. Genome Biol. 15, 550 (2014).

47. Yu, G., Wang, L.-G., Han, Y. & He, Q.-Y. clusterProfiler: an R package for comparing biological themes among gene clusters. OMICS 16, 284–287 (2012).

48. Chen, E. Y. et al. Enrichr: interactive and collaborative HTML5 gene list enrichment analysis tool. BMC Bioinformatics 14, 128 (2013).

49. Wickham, H. ggplot2: Elegant Graphics for Data Analysis. (Springer International Publishing, 2016).

50. The survival package in R. Design of Experiments for Reliability Achievement 339–350 Preprint at https://doi.org/10.1002/9781119237754.app1 (2022).

51. Kassambara, Kosinski, Biecek & Fabian. survminer: Drawing Survival Curves using ‘ggplot2’. R package version 0.3.

52. Hastie, T., Qian, J. & Tay, K. An Introduction to glmnet. CRAN R Repositary (2021).

53. Sing, T., Sander, O., Beerenwinkel, N. & Lengauer, T. ROCR: visualizing classifier performance in R. Bioinformatics 21, 3940–3941 (2005).

54. Yu, G., Wang, L.-G. & He, Q.-Y. ChIPseeker: an R/Bioconductor package for ChIP peak annotation, comparison and visualization. Bioinformatics 31, 2382–2383 (2015).

55. pheatmap: Pretty Heatmaps. Comprehensive R Archive Network (CRAN) https://cran.r-project.org/web/packages/pheatmap/index.html.

56. Thorvaldsdottir, H., Robinson, J. T. & Mesirov, J. P. Integrative Genomics Viewer (IGV):high-performance genomics data visualization and exploration. Briefings in Bioinformatics vol. 14 178–192 Preprint at https://doi.org/10.1093/bib/bbs017 (2013).

