## Supplementary Figures 1-13 for "Genomic analysis reveals HDAC1 regulates clinically relevant transcriptional programs in pancreatic cancer"

**a**

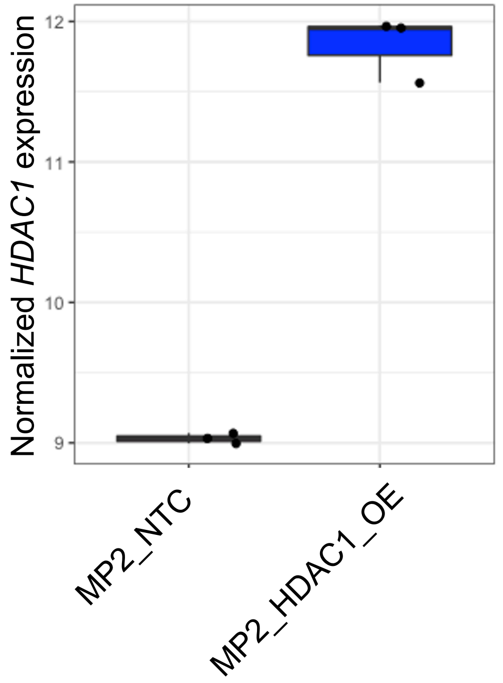


**b**

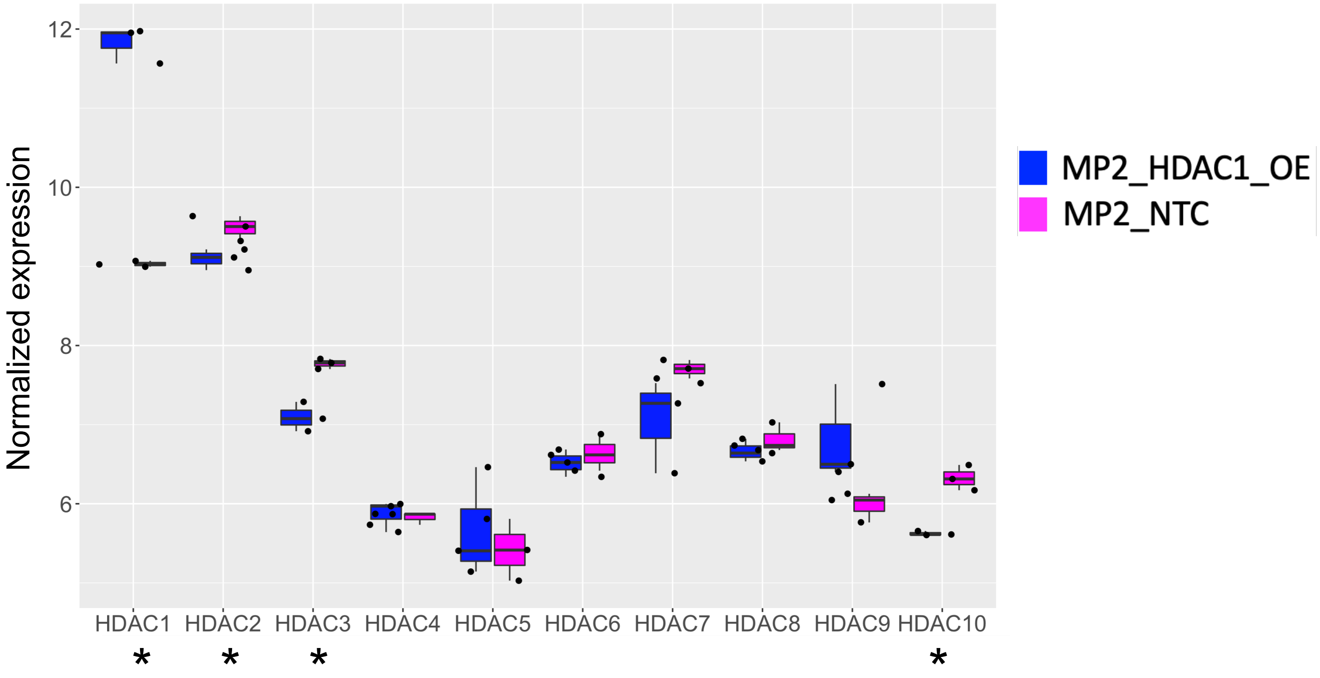


**Supplementary Fig. 1. Normalized gene expression of HDACs.**

**a)** Boxplot of normalized gene expression of *HDAC1* in MP2_HDAC1_OE (blue) and MP2_NTC (pink) cell lines. P-values were calculated using an unpaired two-tailed t-test, p = 3.124e-05. **b)** Boxplot of normalized gene expression data of HDACs for MP2_HDAC1_OE (blue) and MP2_NTC (pink) cell lines. Bar = median. P-values were calculated using an unpaired two-tailed t-test, *p < 0.05.

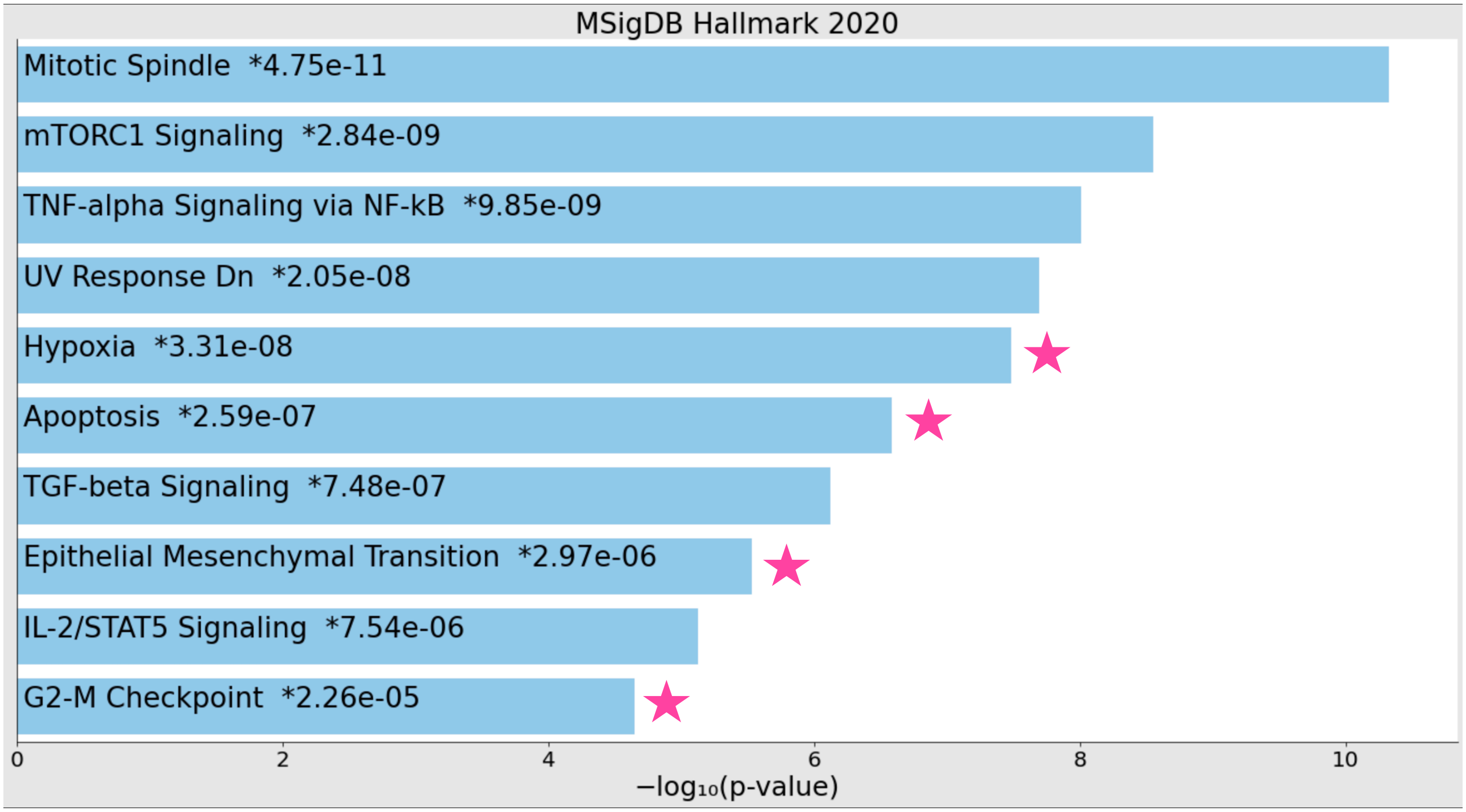


**Supplementary Fig. 2. Analysis of differentially expressed genes revealed pathways altered by *HDAC1* overexpression.**
Barchart showing the top 10 enriched terms from pathway enrichment analysis using genes differentially expressed upon *HDAC1* overexpression. The p-value is shown next to each term. Stars indicate pathways noted in text.

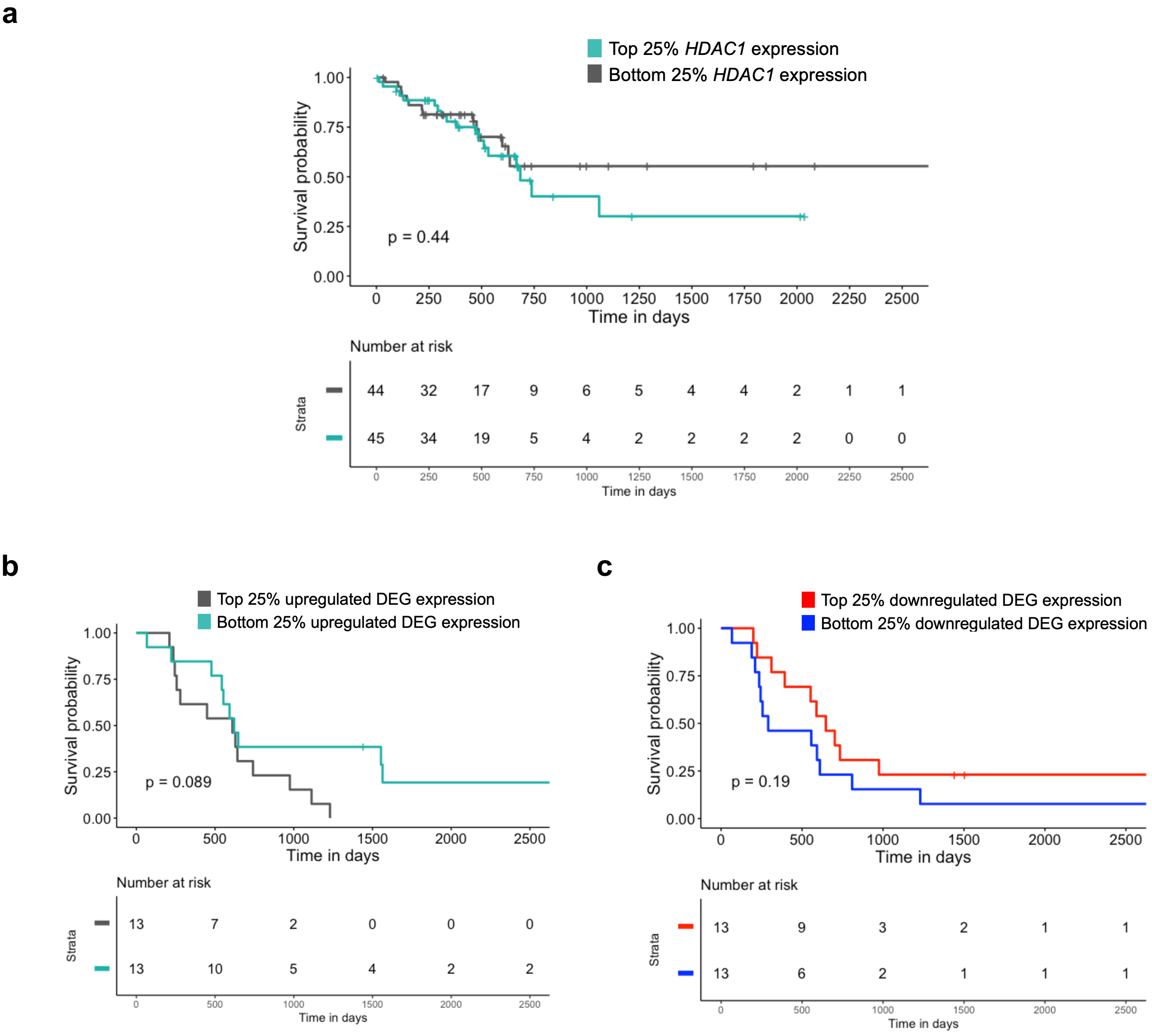


**Supplementary Fig. 3. Overall survival of PDAC patients grouped by *HDAC1* expression.**
**a)** Overall survival of TCCA PDAC patients (n = 90) with top and bottom 25% of *HDAC1* expression. P-value was derived using log-rank test. **b)** Overall survival of an independent cohort of PDAC patients (n = 26) with top (grey) and bottom (teal) 25% of average gene expression of upregulated genes (n = 216). P-values were derived using log-rank test. Only 212 of 216 genes measured in the TCGA cohort were available in this independent cohort, therefore only 208 genes used in this survival analysis. P-values were derived using log-rank test. **c)** Overall survival of an independent cohort of PDAC patients (n = 26) with top (red) and bottom (blue) 25% of average gene expression of downregulated genes (n = 100). Only 100 of 106 genes measured in the TCGA cohort were available in this independent cohort, therefore only 100 genes used in this survival analysis. P-values were derived using log-rank test.

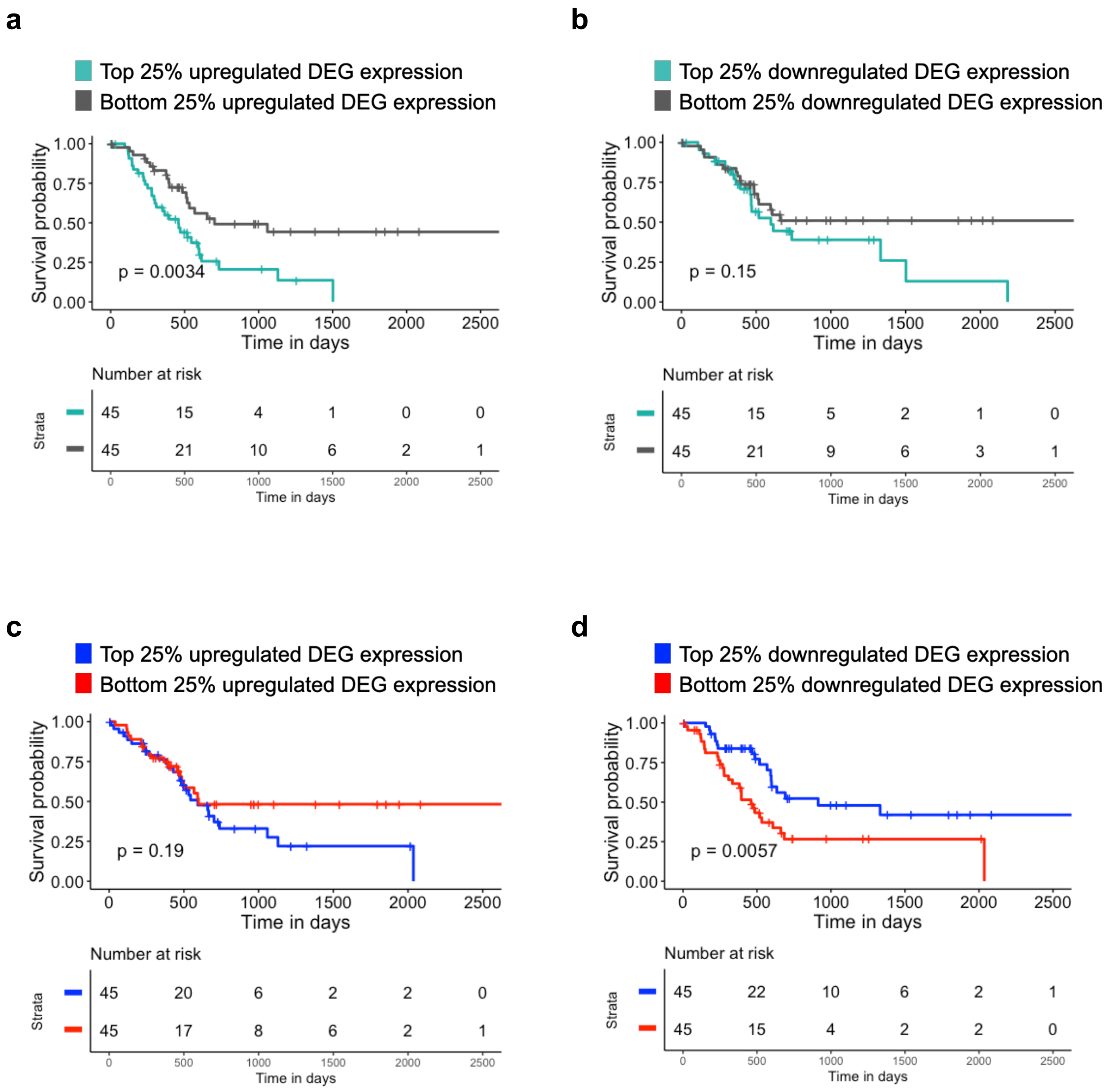


**Supplementary Fig. 4. Survival analysis of the top 322 DEG in cell lines and TCGA PDAC tumors.** Overall survival of TCGA PDAC patients (n = 90) plotted using the top 322 differentially expressed genes (DEG) in MP2_HDAC1_OE cell lines. DEG were ranked by adjusted p-value. 63 DEG were not present in TCGA PDAC expression data, therefore the top 322 available of the top 386 DEG were used. The patients with the top (green) and bottom (grey) 25% of mean gene expression of **a)** upregulated DEG (n = 231) and **b)** downregulated DEG (n = 91) were plotted. P-values were derived using log-rank test. Overall survival of TCGA PDAC patients (n = 90) plotted using the top 322 differentially expressed genes (DEG) in TCGA PDAC tumors with high *HDAC1* expression. DEG were ranked by adjusted p-value. The patients with the top (blue) and bottom (red) 25% of mean gene expression of **c)** upregulated DEG (n = 27) and **d)** downregulated DEG (n = 295) were plotted. P-values were derived using log-rank test.

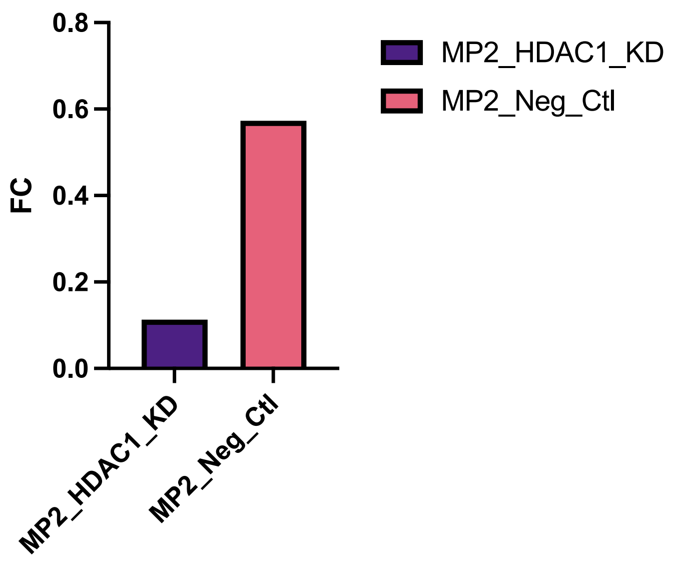


**Supplementary Fig. 5. Quantification of *HDAC1* knockdown with DsiRNA in MP2 cells.**
Boxplot of relative fold change of *HDAC1* in MP2_HDAC1_KD (purple) and MP2_Neg_Ctl (pink) cell lines. Fold change represents relative expression compared to MP2_Neg_Ctl. Values are average of triplicates.

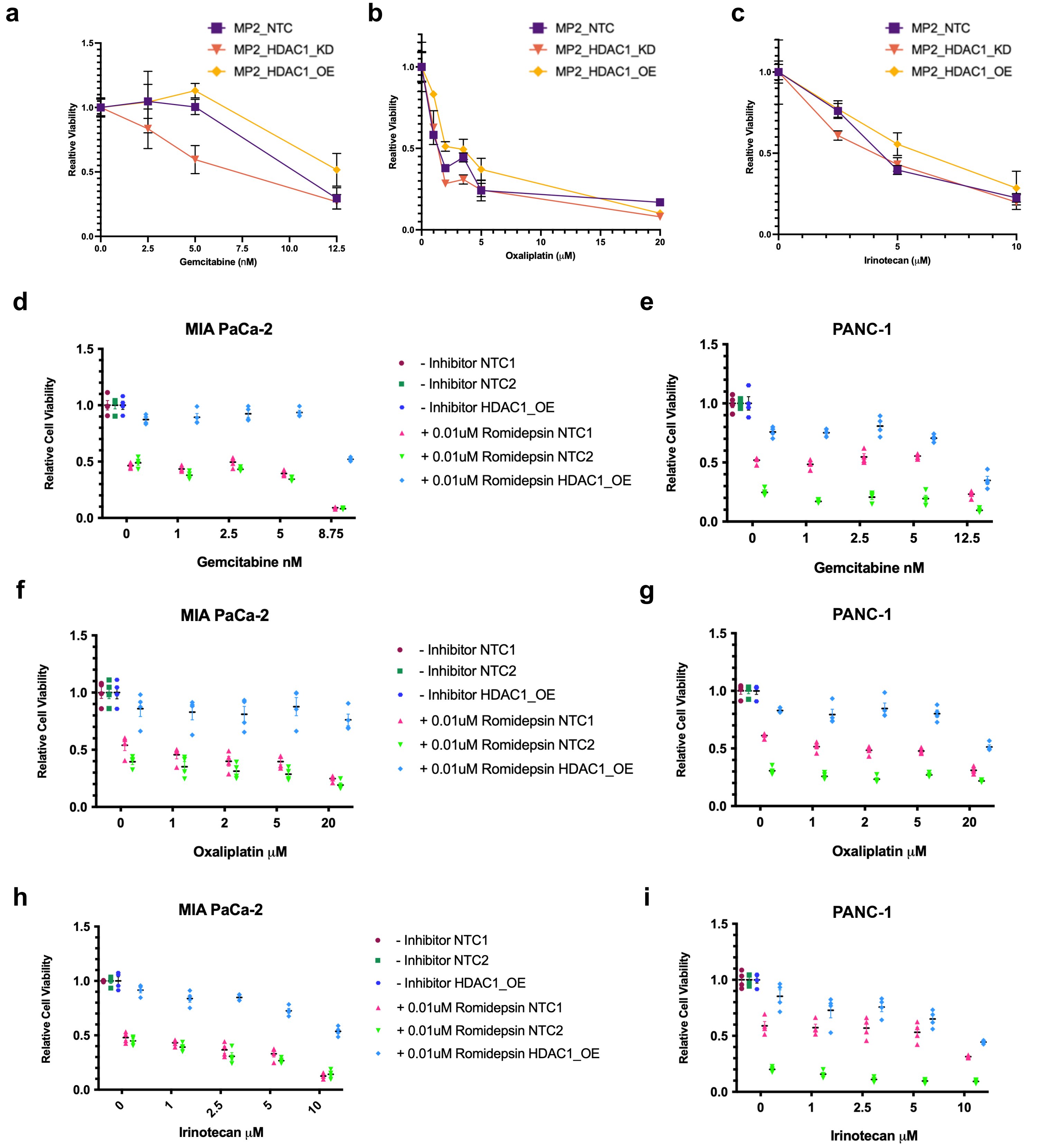


**Supplementary Fig. 6. Quantification of cell viability in PDAC cell lines following treatment of chemotherapeutics.**

**a-c)** Quantification of viability following treatment with **a)** gemcitabine, **b)** oxaliplatin, and **c)** irinotecan in MP2_HDAC1_OE (yellow triangle), MP2_NTC (purple square), and MP2_HDAC1_KD (orange triangle) The bar represents the median. **d-i)** Quantification of viability following treatment in either MIA PaCa-2 or PANC-1 cells with **d-e)** gemcitabine, **f-g)** oxaliplatin, and **h-i)** irinotecan in MP2_HDAC1_OE (blue circles/blue diamond), MP2_NTC1 (burgundy circle/red triangle), and MP2_NTC2 (green square/green triangle) cell lines without or with 0.01μM romidepsin. The bar represents the median.

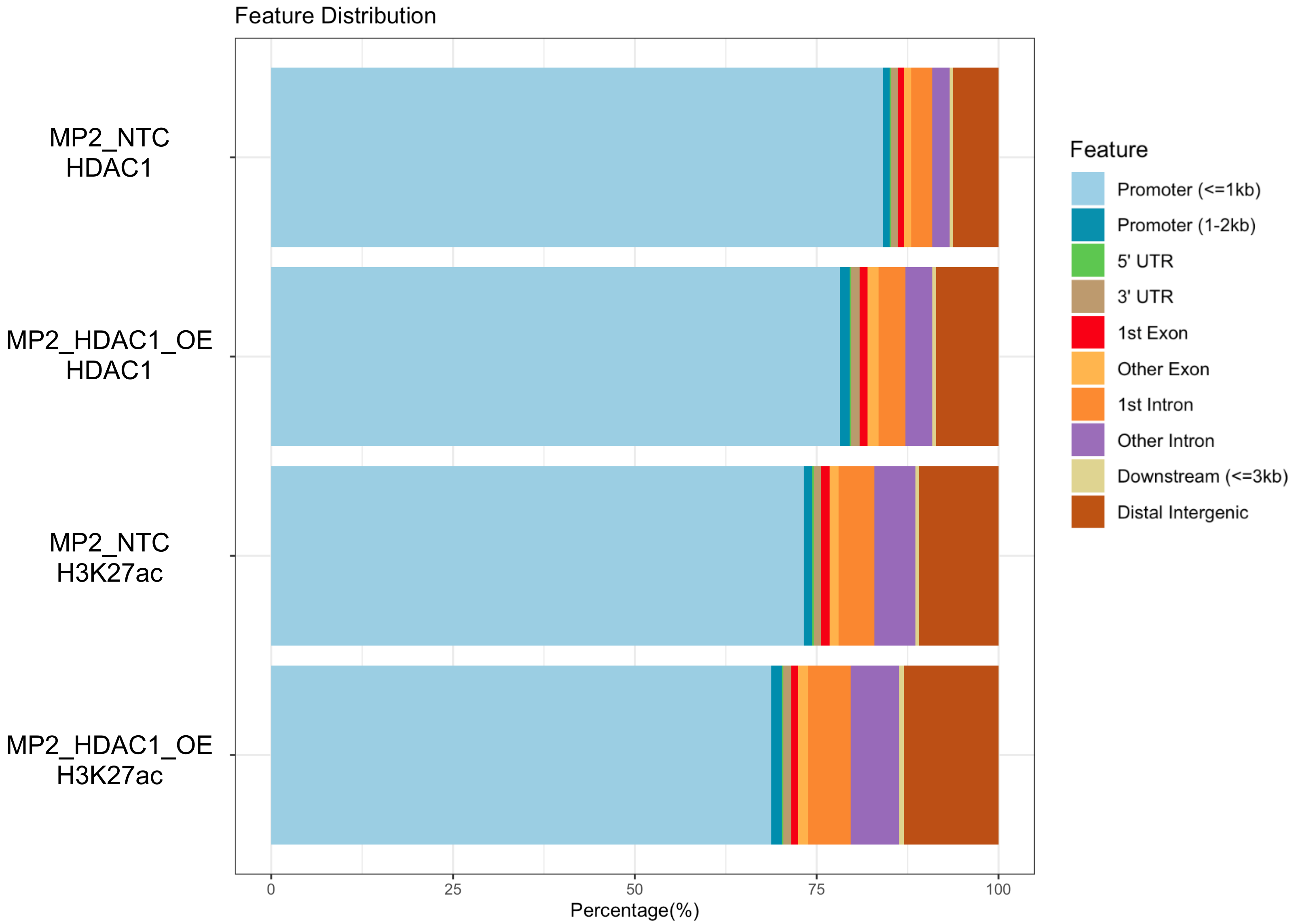


**Supplementary Fig. 7. Feature distribution of ChIP-sequencing peaks.**
Annotation of IDR peaks to genomic regions for H3K27ac in the MP2_HDAC1_OE and MP2_NTC cell lines and HDAC1 IDR peaks in the MP2_HDAC1_OE and MP2_NTC cell lines. Bar graphs were generated using the R package “ChIPseeker”.

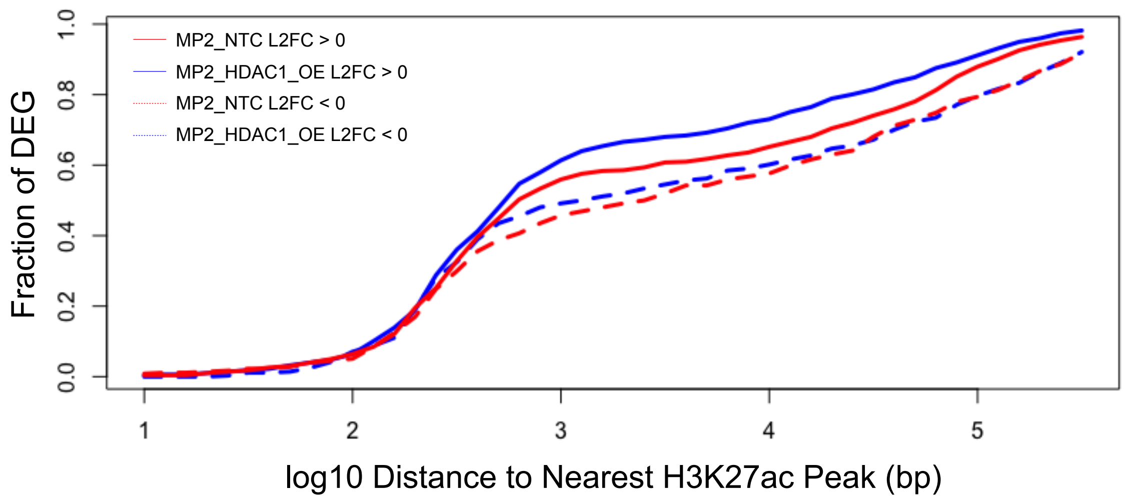


**Supplementary Fig. 8. H3K27 acetylation is more abundant near upregulated genes altered by *HDAC1* overexpression.**

Cumulative distribution graph shows that H3K27ac is more frequently found nearby upregulated DEG. P-values were generated using a Kolmogorov-Smirnov test. MP2_HDAC1_OE_UP (solid blue line) vs MP2_HDAC1_OE_Down (dashed blue line), p = 0.08739. MP2_NTC_UP (solid red line) vs MP2_NTC_Down (dashed red line), p = 0.3421. Up and down regulated genes were chosen from analysis using DESeq2, Log2FoldChange > 0 and Log2FoldChange < 0, respectively.

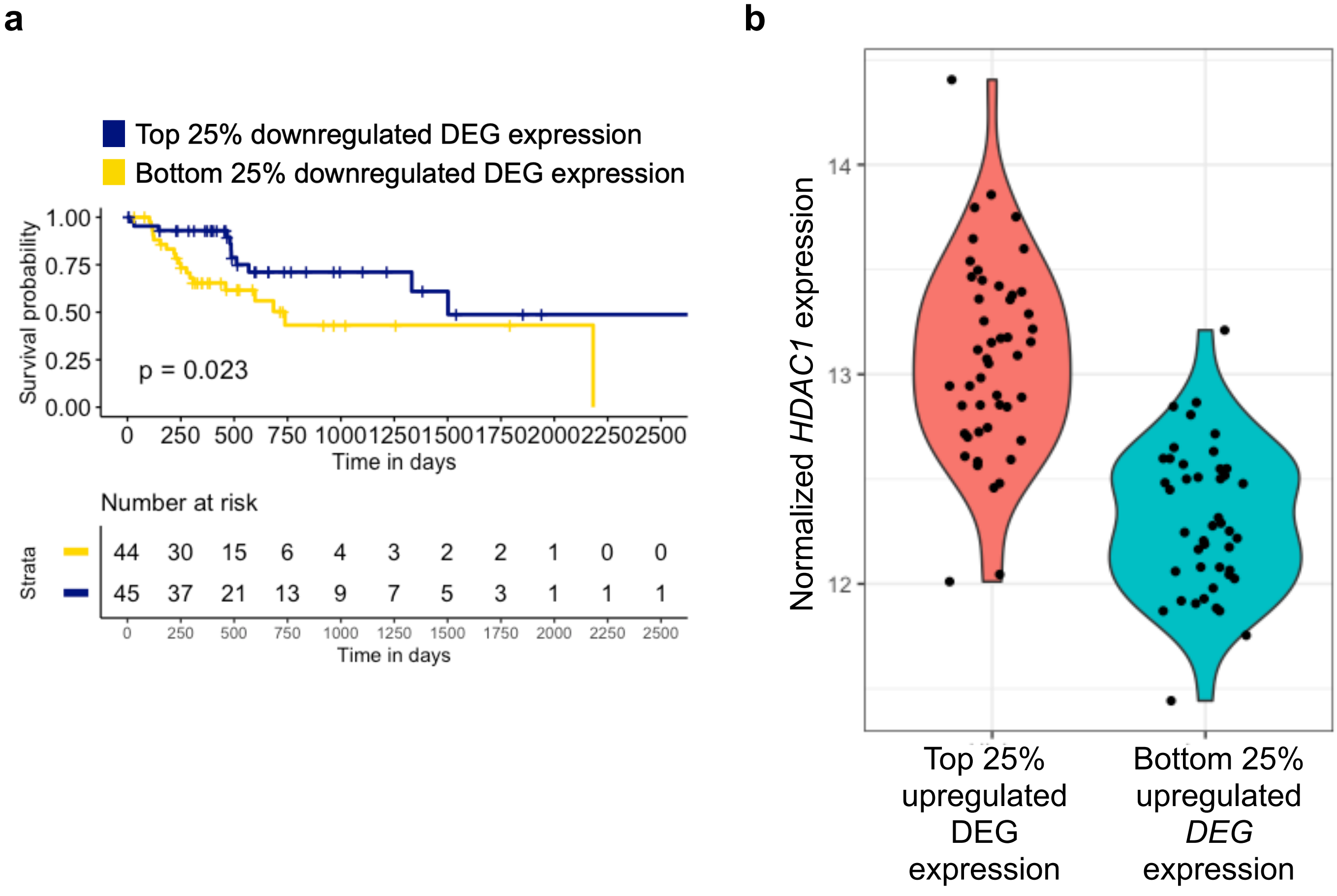


**Supplementary Fig. 9. Analysis of DEG with increased HDAC1 binding and H3K27 acetylation in their promoter.**

**a)** Overall survival of TCCA PDAC patients (n = 90) with top (navy) and bottom (yellow) 25% of average gene expression of downregulated DEG with increased HDAC1 binding and H3K27 acetylation in their promoter upon *HDAC1* overexpression. P-value was derived using log-rank test.

**b)** Violin plot showing normalized *HDAC1* expression for the TCGA PDAC patients in the top 25% (salmon) and bottom 25% (teal) of average upregulated DEG expression. P-value was calculated using an unpaired two-tailed t-test, p = 3.869e-14**.**

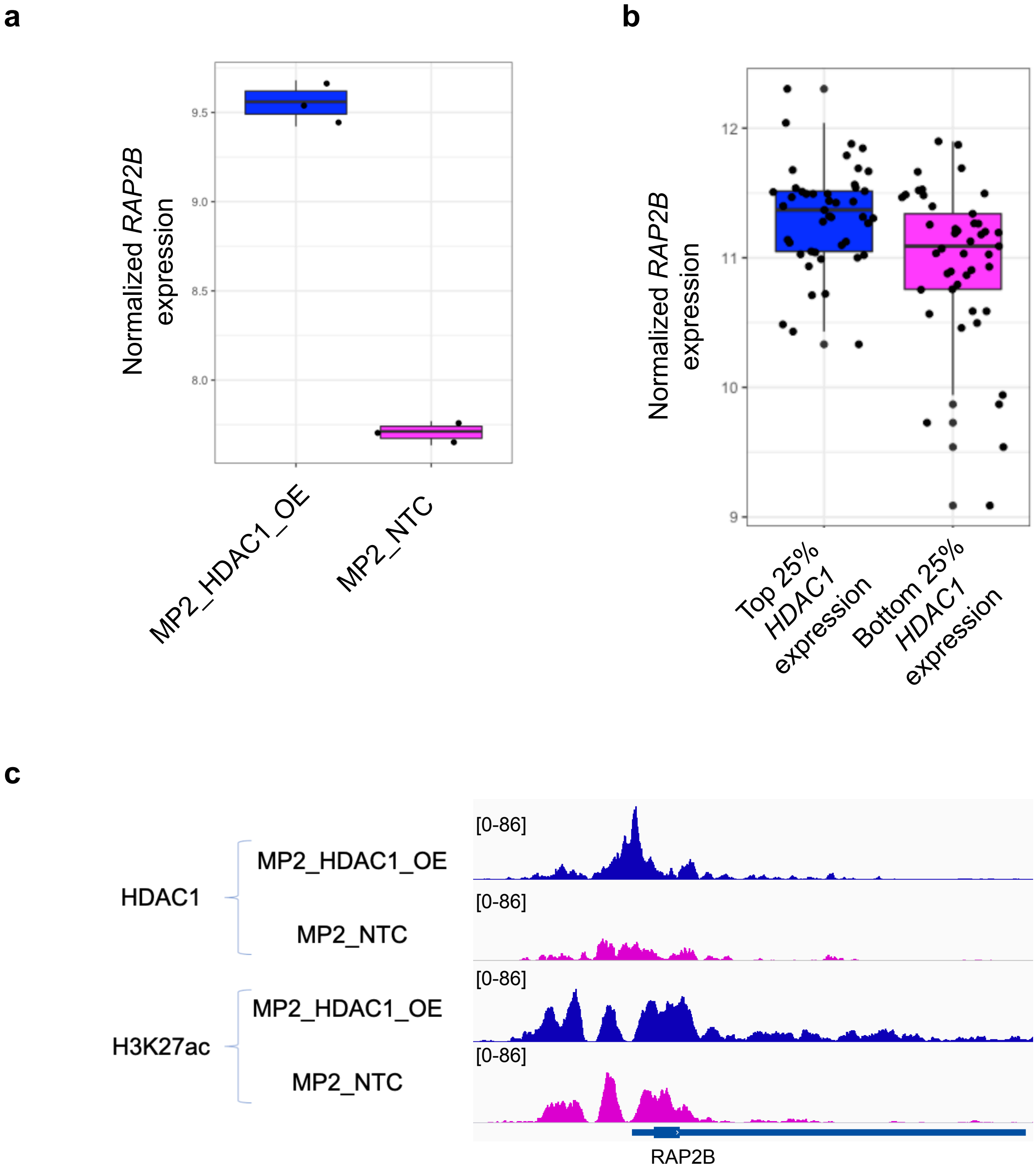


**Supplementary Fig. 10. RAP2B activity upon *HDAC1* overexpression in PDAC cell lines and TCGA PDAC tumors.**
**a)** Normalized expression of *RAP2B* in MP2_HDAC1_OE (blue) and MP2_NTC (pink) cell lines. The bar is the median. P-value was calculated using an unpaired two-tailed t-test, p = 2.594e-05. **b)** Normalized expression of *RAP2B* in the TCGA PDAC samples with the top 25% *HDAC1* expression (blue, n = 45) and bottom 25% *HDAC1* expression (pink, n = 45). The bar is the median. P-value was calculated using an unpaired two-tailed t-test, p = 0.002223. **c)** ChIP-seq analysis of the RAP2B promoter showing increased HDAC1 and H3K27ac peak height in MP2_HDAC1_OE (blue) cells compared to MP2_NTC (pink) cells.

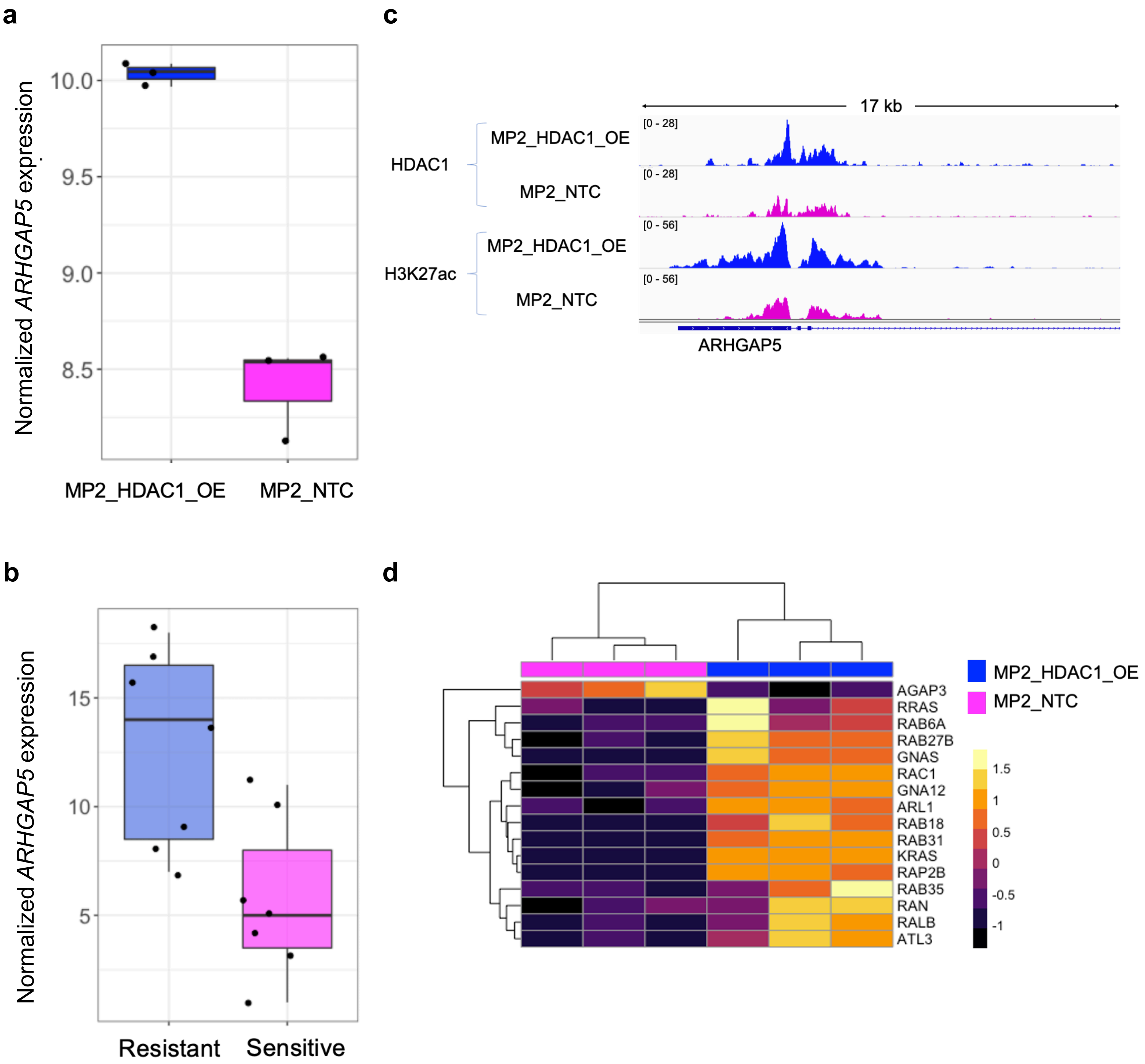


**Supplementary Fig. 11. HDAC1 overexpression is associated with increased GTPase activity.**
**a)** Normalized expression of *ARHGAP5* in MP2_HDAC1_OE (blue) and MP2_NTC (pink) cell lines. The bar is the median. P-values were calculated using an unpaired two-tailed t-test, p = 0.0003479. **b)** Normalized expression of *ARHGAP5* in PDAC cell lines resistant (blue) and sensitive (pink) to gemcitabine. The bar is the median. P-values were calculated using an unpaired two-tailed t-test, p = 0.00831. **c)** ChIP-seq analysis of the ARHGAP5 promoter showing increased HDAC1 and H3K27ac peak height in MP2_HDAC1_OE (blue) cells compared to MP2_NTC (pink) cells. **d)** Expression of DEG involved in GTPase activity in MP2_HDAC1_OE (blue) and MP2_NTC (pink) cell lines. Each column represents a replicate of the noted cell line. The color scale denotes the z-score of each gene.

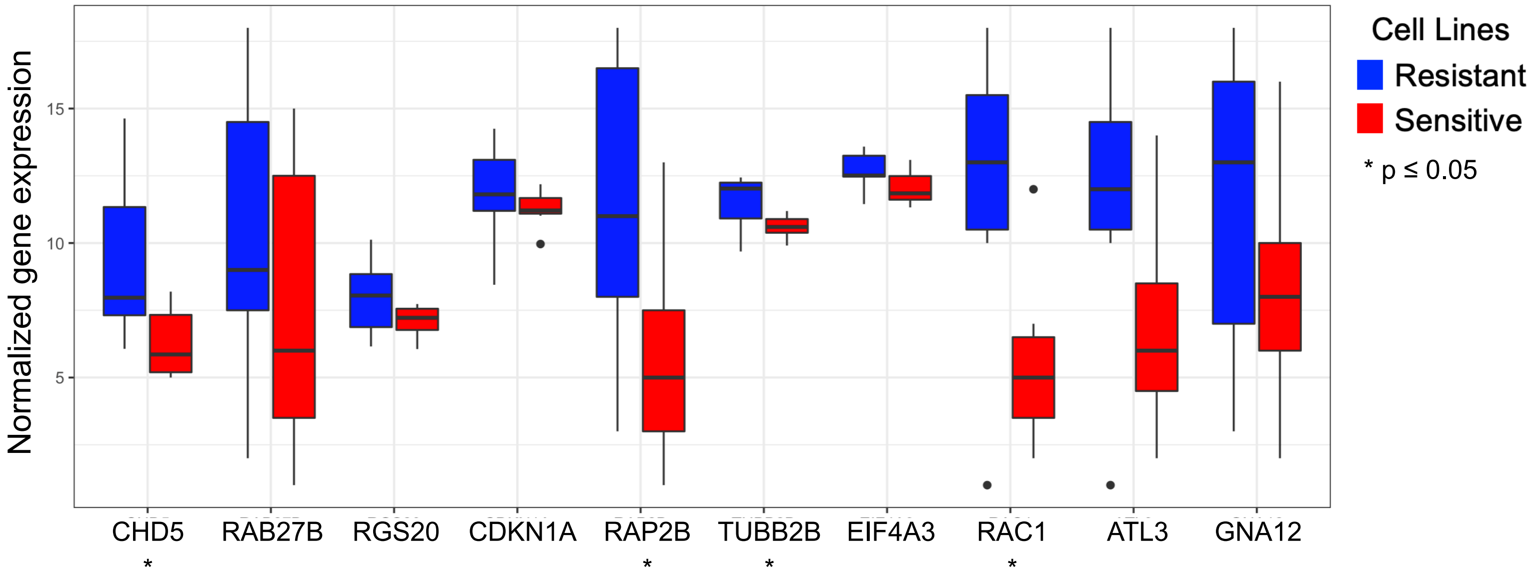


**Supplementary Fig. 12. Normalized expression of genes involved in GTPase activity in chemo resistant and sensitive PDAC cell lines.**
Boxplots comparing normalized gene expression of the top 10 upregulated genes that are upregulated upon *HDAC1* overexpression and enriched for GO terms involved in GTPase activity. The boxplots include 14 PDAC cell lines resistant (n = 7, blue) or sensitive (n = 7, red) to gemcitabine. Bars represent the median. P-values were calculated using an unpaired two-tailed t-test, *p ≤ 0.05.

a

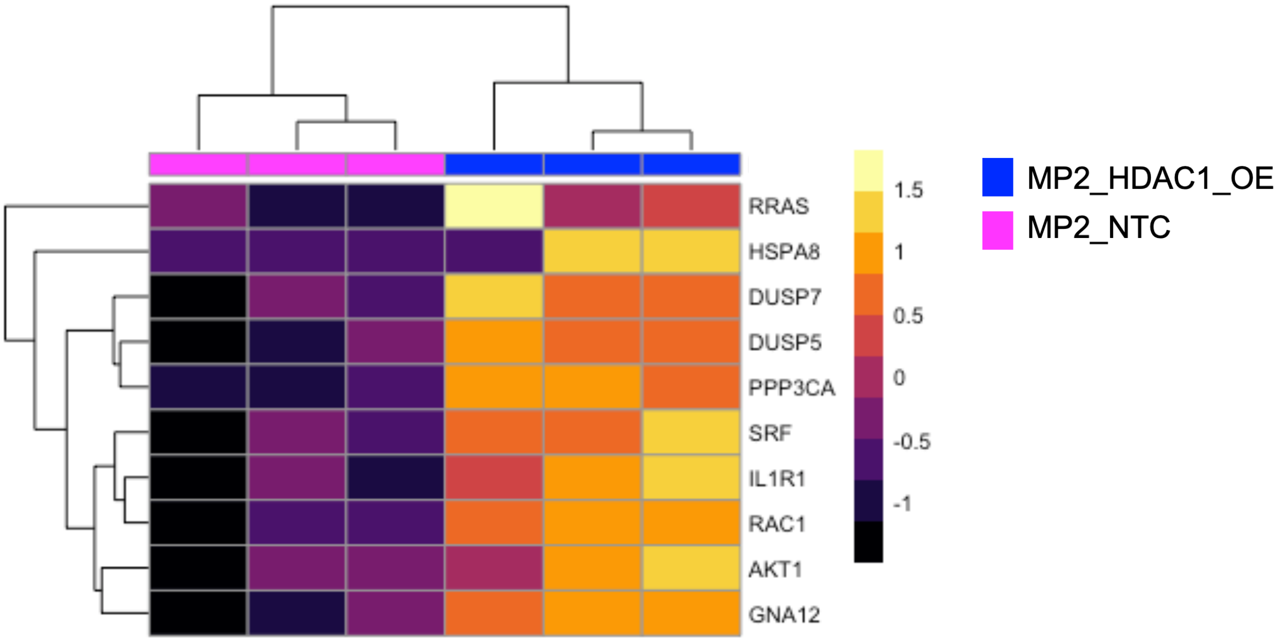


b

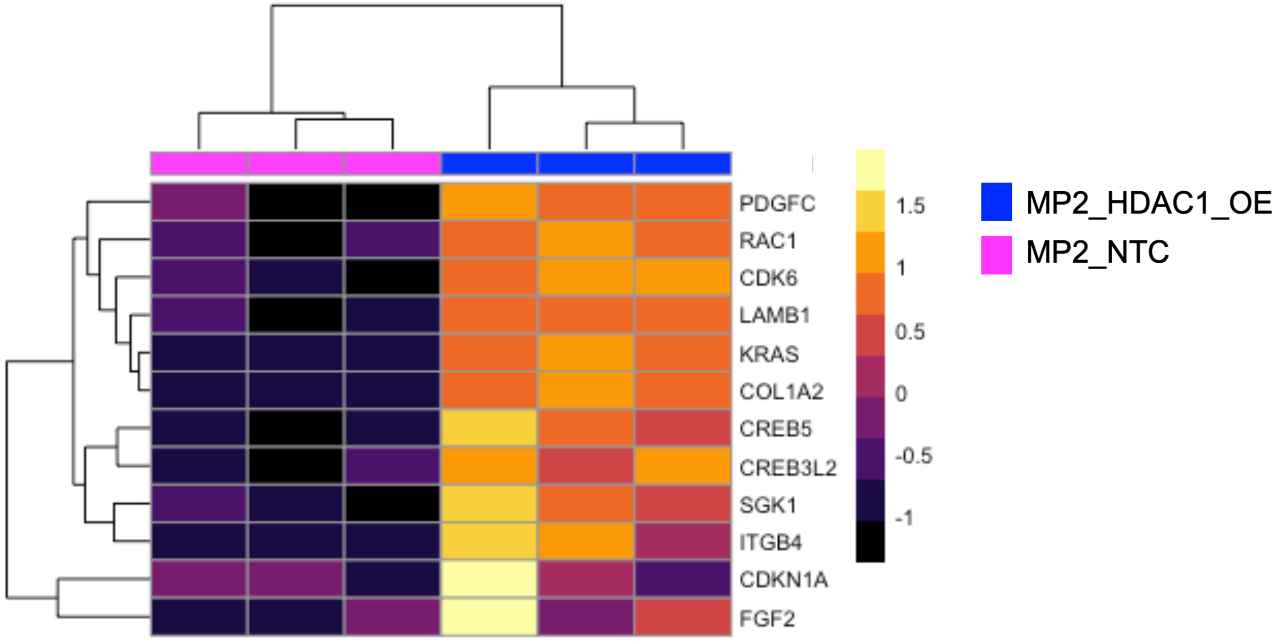


**Supplementary Fig. 13. Expression of genes in MAPK and PI3K pathway are increased upon *HDAC1* overexpression.**
Expression of genes in MP2_HDAC1_OE and MP2_NTC cell lines that are involved in two pathways **(a)** MAPK and **(b)** PI3K. Each column represents a replicate of the denoted cell line. The color scale denotes the z-score of each gene.
